# Does land use matter? Carbon consequences of alternative land use futures in New England

**DOI:** 10.1101/2021.01.08.425951

**Authors:** Meghan Graham MacLean, Matthew Duveneck, Joshua Plisinski, Luca Morreale, Danelle Laflower, Jonathan Thompson

## Abstract

Globally, forests play an important role in climate change mitigation. However, land-use impacts the ability of forests to sequester and store carbon. Here we quantify the impacts of five divergent future land-use scenarios on aboveground forest carbon stocks and fluxes throughout New England. These scenarios, four co-designed with stakeholders from throughout the region and the fifth a continuation of recent trends in land use, were simulated by coupling a land cover change model with a mechanistic forest growth model to produce estimates of aboveground carbon over 50 years. Future carbon removed through harvesting and development was tracked using a standard carbon accounting methodology, modified to fit our modeling framework. Of the simulated changes in land use, changes in harvesting had the most profound and immediate impacts on carbon stocks and fluxes. In one of the future land-use scenarios including a rapid expansion of harvesting for biomass energy, this changed New England’s forests from a net carbon sink to a net carbon source in 2060. Also in these simulations, relatively small reductions in harvest intensities (e.g., 10% reduction), coupled with an increased percent of wood going into longer-term storage, led to substantial reductions in net carbon emissions (909 MMtCO_2_eq) as compared to a continuation of recent trends in land use. However, these projected gains in carbon storage and reduction in emissions from less intense harvesting regimes can only be realized if it is paired with a reduction in the consumption of the timber products, and their replacements, that otherwise would result in additional emissions from leakage and substitution.

## INTRODUCTION

Forest carbon plays a key role in regulating the climate system (Houghton et al. 2012, Williams et al. 2012, Reinmann et al. 2016, Ma et al. 2020, Finzi et al. 2020). Forest land use, including timber harvest and conversion for developed uses, has significant impacts on forest carbon dynamics and, thus, future land use has the potential to mitigate or exacerbate climate change (Pan et al. 2011, Butler et al. 2015, Woodall et al. 2015, Le Quéré et al. 2018). Mechanistic models of forest carbon dynamics, coupled to simulations of co-designed land-use scenarios, offer a robust approach to identifying and planning for sustainable land-use pathways.

Like much of the global temperate forest biome, the northeastern U.S. has significant capacity to increase its forest carbon stocks through natural regrowth (Cook-Patton et al. 2020a). Continued forest growth and recovery from Colonial-era land use remains the most significant driver of aboveground carbon dynamics in this region (Thompson et al. 2013, Puhlick et al. 2017, Duveneck et al. 2017). However, the ability of the region to continue to serve as a carbon sink is threatened by the current land-use regime. Since the 1980s, land-use and land-cover (LULC) change, particularly the expansion of low-density residential development, has resulted in the net loss of approximately 387,000 ha of forest cover across the six New England states (Olofsson et al. 2016), reducing stocks and the capacity for future terrestrial carbon sequestration (Reinmann et al. 2016, Thompson et al. 2017b). If rates and spatial patterns of forest conversion continue as they have from 1990-2010 through 2050, an additional 0.5 million ha of forest land could be lost to development with consequential impacts to carbon storage and sequestration (Thompson et al. 2017b). Even more importantly, despite recent reductions in timber harvesting throughout much of southern New England (Kittredge et al. 2017), harvesting remains the primary driver of mature tree mortality and carbon loss throughout the region (Canham et al. 2013, Harris et al. 2016, Thompson et al. 2017a, Ma et al. 2020). Therefore, it is important to understand how changes in future land-use patterns, including both development and harvesting, affect the total carbon storage in New England’s forests and elsewhere.

Understanding the carbon impacts of future land-use choices in a heavily forested and heavily populated region, such as New England, can help guide future policy and land use, but anticipating the future conditions of regional ecosystems where small private landowners dominate is challenging. Sixty-five percent of New England forests are owned and managed by more than 200,000 family forest owners, each making land-use decisions based on their own priorities (Butler et al. 2016). The sum of these choices has significant impacts on the carbon storage potential of New England forests. Given that predicting the future of these socio-ecological systems is impossible, analyzing alternative land-use scenarios offers a robust way to plan for the future (McBride et al. 2017, 2019). Land-use scenarios describe potential future socio-ecological dynamics and their consequences, using internally consistent assumptions about major drivers of change (Li et al. 2008, Schulp et al. 2008, Sleeter et al. 2012, Popp et al. 2014). Increasingly, scenarios are co-designed with stakeholders who, through a structured process, collectively envision possible future land-use pathways (Bradfield et al. 2005, McBride et al. 2017).

In this analysis we evaluate the consequences of five land-use scenarios for forest carbon in New England. One scenario represents a linear continuation of the recent trends in land use, including land-cover change and harvesting (Duveneck and Thompson 2019), and four divergent, alternative scenarios that were co-designed by more than 150 stakeholders (e.g., conservationists, planners, resource managers, landowners, and scientists) as part of the New England Landscape Futures (NELF) project (Figure 1). The scenario co-design process was described in detail by McBride et al. (2017) and the process of translating the qualitative scenarios into simulations of land-cover change was described by Thompson et al. (2020). The described NELF alternative scenarios are highly divergent in terms of the types, intensities, and spatial allocation of land use and, thus, represent a wide range of potential futures for the region’s forests and the services they provide (Figure 2). The land-cover change simulations have subsequently been used to evaluate a range of future outcomes, including flood potential (Guswa et al. 2020), conservation priorities (*Losing Ground: Nature’s Value in a Changing Climate, Sixth Edition of the Losing Ground series 2020*, Thompson et al. 2020), and wildlife habitat (Pearman-Gillman et al. 2020a, 2020b).

**Figure 1.**
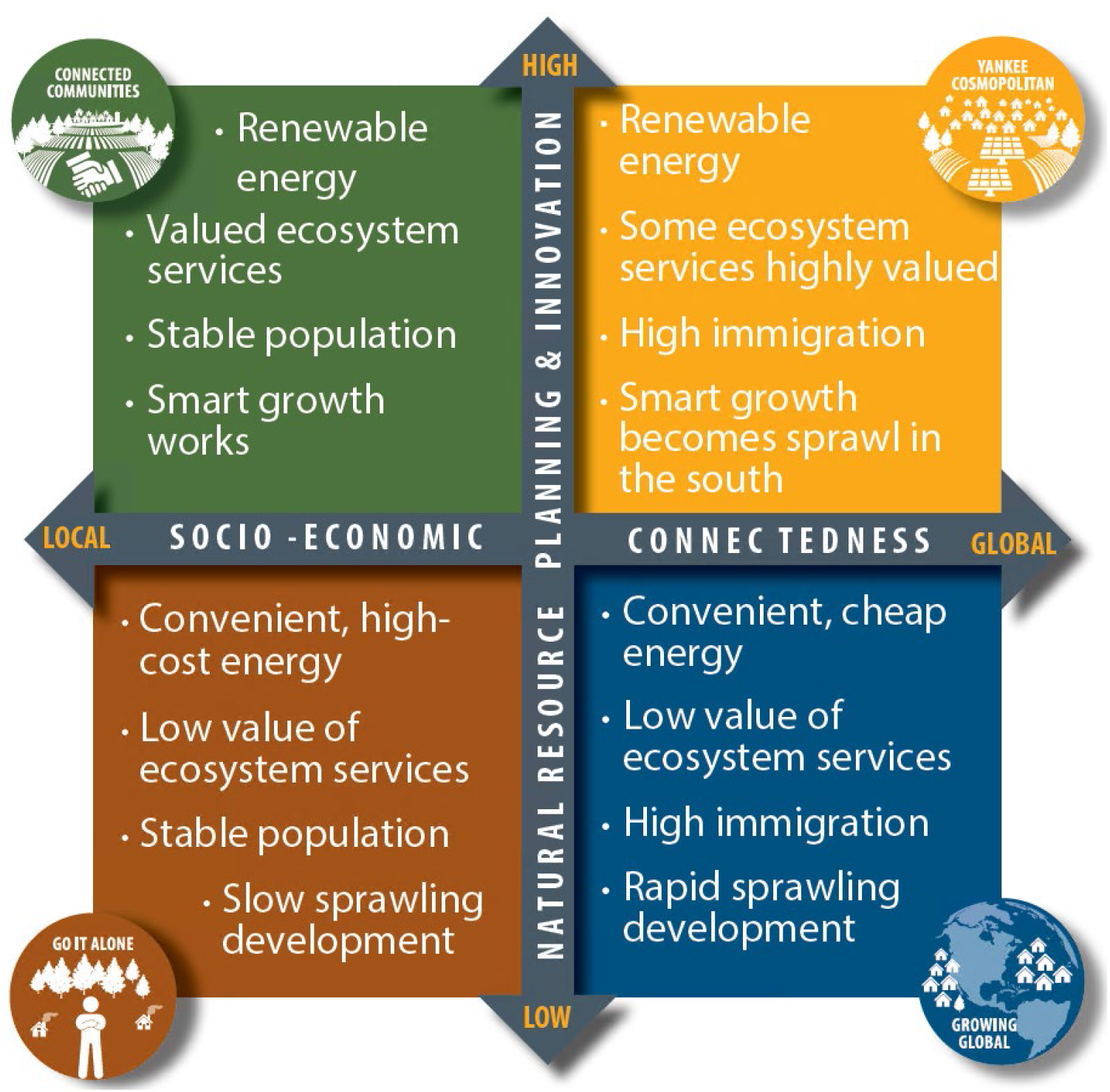
New England Landscape Futures (NELF) scenarios. The four scenarios were articulated along two axes that were identified as the two drivers of greatest influence and uncertainty for future land-use change.

**Figure 2.**
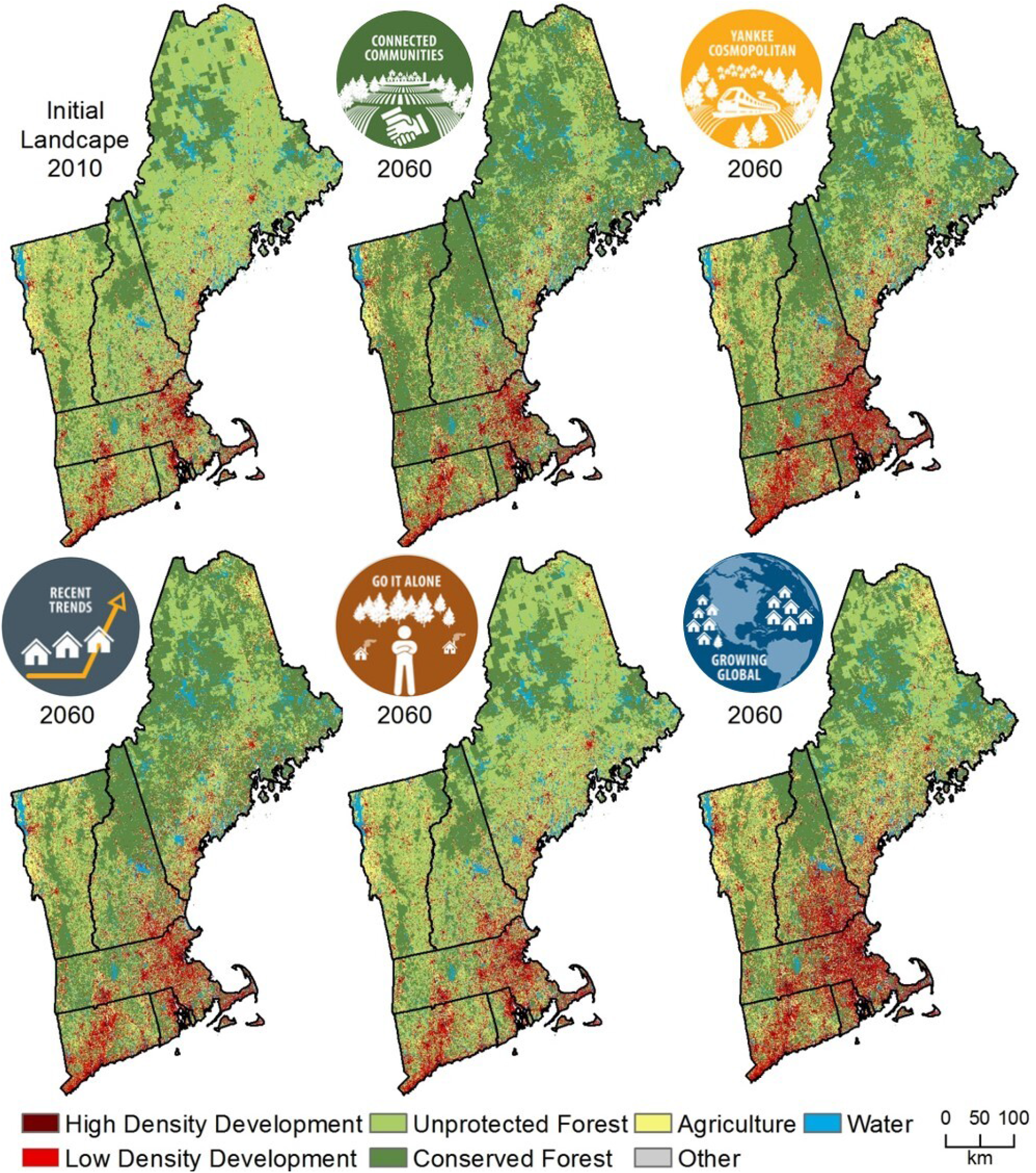
The modeled land-cover change of recent trends in land-cover change as well as the four NELF stakeholder scenarios.

Previously, we evaluated the impacts of a continuation of recent trends in harvesting and development on New England forests (Duveneck and Thompson 2019). This scenario assumed a continuation of the patterns of land use, including development and timber harvesting, observed over the last several decades. Recent trends in development patterns project an increase in development in the southern metropolitan areas as northern rural areas become less populated (Thompson et al. 2020) (Figure 2). Under these assumptions, land use reduced carbon storage by 16% over fifty years, as compared to a counterfactual scenario with no land use (i.e., no development or harvesting). Ownership patterns, from small family forest owners to large industrial timberlands, explained a large part of the landscape variation in carbon dynamics (Duveneck and Thompson 2019), highlighting the importance of landowner impacts on carbon due to the disjointed management decisions of many private landowners. In contrast, climate change alone increased carbon stocks by only 8% in this recent trends scenario, due in large part to longer growing seasons (Duveneck et al. 2017).

Here we expand and improve our previous analysis to include the four co-designed scenarios and a more in-depth estimation of the changes in forest carbon due to future land use. These four co-designed scenarios present a range of future land-use regimes, in terms of development and harvesting, that impact future carbon storage and emissions, and therefore elucidate how changes in land-use can influence the total carbon balance of New England’s forests. We also use an improved calibration and validation scheme to evaluate aboveground carbon accumulation, and we include a more complete accounting of the carbon dynamics that includes the removed aboveground carbon in all of our future land use scenarios (Smith et al. 2006, Reinmann et al. 2016, Ma et al. 2020). Specifically, we ask: how do characteristics of the NELF scenarios’ envisioned land-use regimes (i.e., harvest intensity and extent, forest loss to development, and wood product innovation) differentially drive changes in future aboveground carbon emissions, storage, and sequestration.

## METHODS

### Study Area

The study area is in the northeastern United States and encompasses the six New England states (Connecticut, Rhode Island, Massachusetts, Vermont, New Hampshire, and Maine) (Figure 3). The region contains approximately 13 million hectares of forest which cover approximately 80% of the land area. Forest types in the region span from oak pine forests in the south, to northern hardwoods across most of the central region, to boreal forests in the north (Duveneck et al. 2015). Likewise, mean annual temperatures span a north-south gradient from 3 to 10 °C. Mean annual precipitation in the region ranges from approximately 79 to 255 cm, with higher rates of precipitation at higher elevation (Huntington et al. 2009). The New England region is inhabited by approximately 15 million people (2018 U.S. Census). Most of the people in New England are concentrated in the metropolitan areas of Southern New England (e.g., Boston, MA; Hartford, CT; and Providence, RI) with much of the rural north sparsely populated. The majority of forest land in the region is owned by private landowners with relatively small parcels (< 10 ha) who are largely uncoordinated in the management of their lands (Butler et al. 2016). Corporate and investment timber lands are concentrated in the north, primarily in Maine.

**Figure 3.**
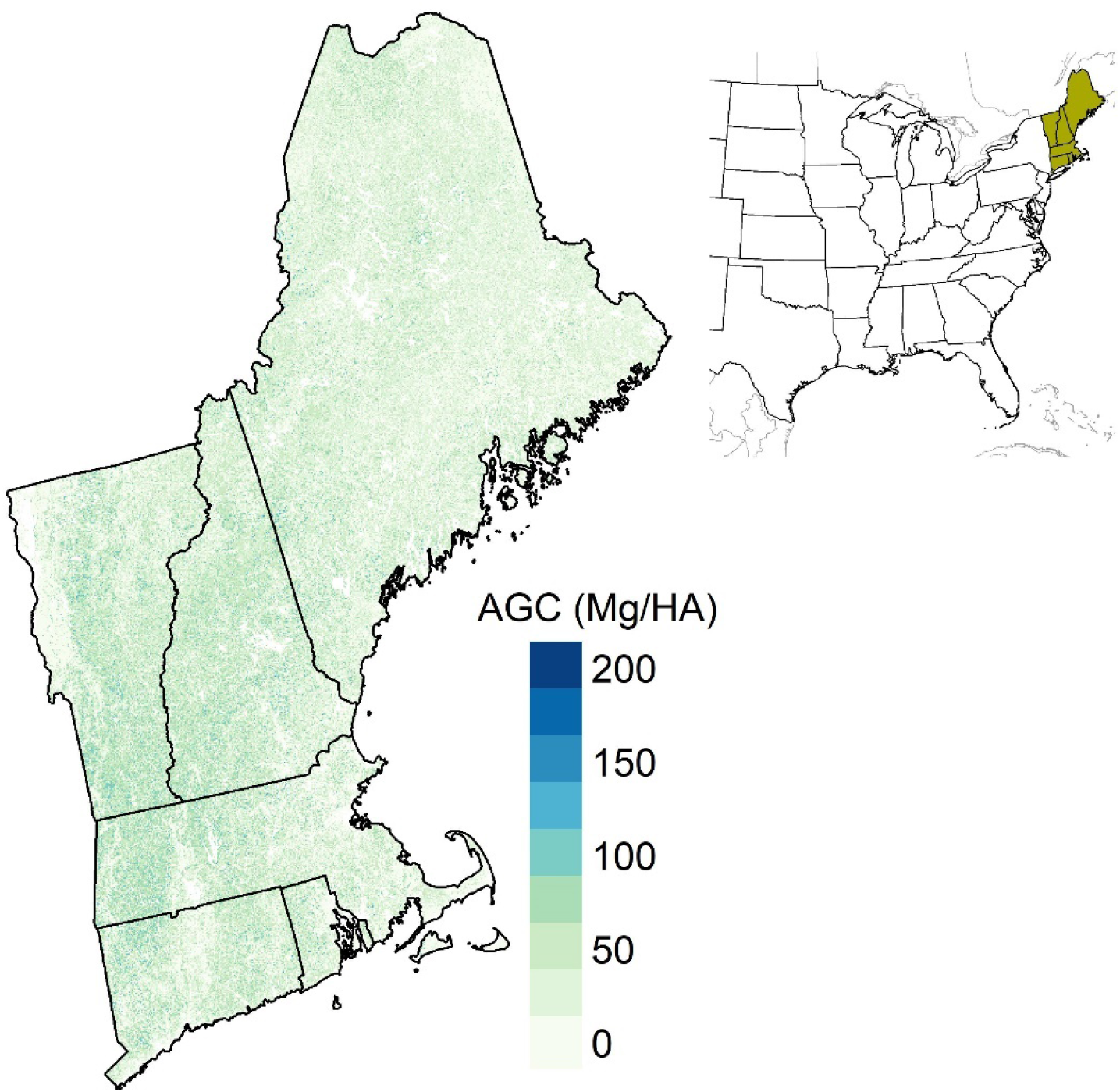
New England study area map showing aboveground carbon (AGC; in Mg ha^−1^) for 2010. Inset map shows study area within eastern United States.

### Modeling framework

We simulated the effects of the five divergent land-use scenarios as described by stakeholders as part of the NELF project (Thompson et al. 2020), on aboveground forest carbon in New England from 2010 to 2060. We used a forest composition raster with 250 m resolution from Duveneck et al. (2015) as our initial forest area, biomass, and composition for 2010 (Figure 3). This initial condition map was based on an imputation of USDA Forest Inventory and Analysis (FIA) plots (Bechtold and Patterson 2005). Belowground carbon, while quite important, was outside the scope of this research. To track aboveground carbon storage and emissions from land use (i.e., development and harvesting), we employed multiple models linked together to form our modeling framework. We first utilized the outputs from the NELF land-cover change simulations modeled using Dinamica – EGO, and described previously in Thompson et al. (2020), to spatially allocate forest land-cover transitions within each scenario (see Appendix I). Within the forested area, we simulated forest growth and succession using LANDIS-II (Mladenoff and He 1999, Scheller et al. 2007) with the PnET-Succession module (de Bruijn et al. 2014) from 2010 to 2060 at 10-year time steps. We simulated timber harvesting using the LANDIS-II extension Biomass-Harvest (Gustafson et al. 2000). We then coupled these models to a common carbon accounting framework to track the fate of carbon removed through various land-use practices (Smith et al. 2006). A more complete description of each model component is below.

#### DEVELOPMENT AND CONSERVATION

As described previously in Thompson et al. (2017b, 2020), we used Dinamica – EGO v.2.4.1 (Soares-Filho et al., 2002), a cellular land cover change model, to simulate land-cover transitions for each of the five land-use scenarios based on the individual scenario narratives and stakeholder input on how rates of land-cover change would be different in the co-designed scenarios from those observed in recent trends (Appendix I). Within the land-cover simulations, transition rates allocation parameters were defined individually for each core-based statistical area (CBSA) as defined by the U.S. Census (www.census.gov; accessed 4/20/2019). For areas that did not fall within Census-defined CBSAs, new regions were defined to model land-cover transitions (Thompson et al. 2020). The modeled land covers included forest, agriculture, water, development, along with the transition of some forests to conserved forests (Figure 2). Land-cover transitions of interest to this project included transitions from forest to agriculture, low-density development, and high-density development, as well as from unconserved to conserved forest. For ease, we will refer to the conversion of forest to other land cover types (except water) generically as ‘development.’ Conservation became an important component of the land use simulations, as some of the simulated conserved forest restricted harvesting, and thus impacted the spatial allocation of harvest (see ‘Harvesting’ below for more detail).

The resulting land cover maps from the Dinamica – EGO simulation had a 30 m spatial resolution and included individual maps of land cover for every 10^th^ year of the 50-year simulations, from 2010 to 2050. The 30 m land cover simulation outputs were resampled to 250 m to match the spatial resolution of our forest composition layer. During the resampling process, if there was only partial forest conversion of a single 250 m cell we calculated the proportion of the 250 m cell that was converted from forest to another land cover and removed the appropriate biomass from the 250 m cell to represent the proportional area converted to other land cover. We did not simulate afforestation in these scenarios (i.e., agriculture transitioning to forest) as these patterns are not prevalent in this landscape and were not included in the narratives of the future scenarios.

#### FOREST GROWTH AND MODELING CALIBRATION

For all forested areas in New England, we simulated forest growth using the PnET-Succession extension (v.3.4) (de Bruijn et al. 2014) of the LANDIS-II (v. 7.0) forest simulation model (Scheller et al. 2007). LANDIS-II is a spatially explicit, mechanistic forest landscape model that simulates forest growth, competition, and dispersion within forest raster cells. Rather than model individual trees, LANDIS-II simulates species-age cohorts which mature and disperse within interacting cells. PnET-Succession simulates photosynthesis, respiration, and mortality based on the PnET Carbon Model (Aber et al. 1995) and has been extensively evaluated and utilized in New England (e.g., Duveneck and Thompson 2017, 2019, Liang et al. 2018, McKenzie et al. 2019) and beyond. One of the strengths of the combination of LANDIS-II and PnET-Succession is that it is a mechanistic model based on first principals of forest growth, and therefore useful in simulating the impacts of changes in land use in novel circumstances, such as with climate change (Gustafson 2013, Duveneck and Thompson 2019). Therefore, we used the Regional Conservation Pathway (RCP) 8.5 emission scenario (Stocker et al. 2013) as projected by the Hadley Global Environment v.2-Earth System Global Circulation Model (GCM), downscaled and obtained from the USGS Geo Data Portal (Stoner et al. 2013) to evaluate the impacts of land use, with climate change, for all scenarios. For each NELF scenario simulation, we used LANDIS-II/PnET-Succession to model growth and senescence of aboveground tree biomass, and therefore track carbon stocks and fluxes, for forested areas at 10-year time steps.

To account for carbon loss to natural disturbance, we simulated a low-frequency wind disturbance regime across all scenarios, because this is the primary background natural disturbance occurring across the region. We used the Base Wind extension (Mladenoff and He 1999) for LANDIS-II to emulate these low-severity wind-based mortality events. Specifically, we simulated a wind rotation period of 400 years with a maximum, mean, and minimum patch size of 400, 20, and 6 hectares, respectively. Within each wind patch, the probability of cohort mortality was based on the cohort age, where cohorts that had reached 85% of their age had a mortality probability of 0.65. Younger cohorts had successively lower mortality probabilities.

To evaluate our PnET-Succession parameterization of growth and carbon accumulation on undisturbed sites, we compared the mean county-level annual forest growth from remeasured FIA subplots (Bechtold and Patterson 2005) with simulated forest growth in each county. Specifically, we aggregated tree biomass from FIA subplots that were > 90% forested, and had at least 2 measurements after the year 2000. In addition, we further selected only the plots that were relatively undisturbed (i.e., plots that had not experienced an identified disturbance, nor increased biomass in the remeasurement period). To calculate observed forest growth at the county level, we first summed the live aboveground tree biomass for each FIA subplot for each remeasurement period. Next, we converted these values to carbon (carbon = 0.5 * biomass) and annualized the carbon accumulation using the number of years between remeasurement periods unique to that plot. We then divided each subplot’s carbon accrual by its forested area (i.e., the area of the subplot multiplied by the percent of the subplot that was forested) to produce annualized changes in carbon density (Mg ha^−1^ yr^−1^). Finally, for counties with greater than 10 such FIA plots, we aggregated subplots within each county and calculated mean and standard deviation of carbon density.

To compare these FIA estimates of forest growth with our LANDIS-II simulations of forest growth, we simulated forest growth across New England, from 2010 to 2020, with no impacts from human development or harvest, using our imputed 2010 forest biomass map for our initial forest conditions. This 10-year evaluation time period approximated two FIA remeasurement periods (most FIA plots are revisited in approximately 5-year intervals). We included the wind disturbance regime described above in our simulation of forest growth, since similar light disturbances were also included in the FIA plot data. We then calculated the mean annual change in simulated aboveground carbon accumulation for each New England county. For each county, we compared the annual carbon accumulation observed within FIA plots to those simulated by LANDIS-II. Most simulated and observed county mean carbon accumulation rates were within 25% of each other, and all LANDIS-II means were within one standard deviation of the FIA means (Figure 4). Additionally, the grand means were not significantly different (*p* < 0.05) and differed by less than 1% (FIA 1.451 Mg ha^−1^ yr^−1^, LANDIS-II 1.455 Mg ha^−-1^ yr^−1^). Given the variability of tree growth both in observed tree growth and in the simulations due to the stochastic processes within LANDIS-II, we were satisfied by the overall level of agreement between the simulated and observed growth in FIA plot data.

**Figure 4.**
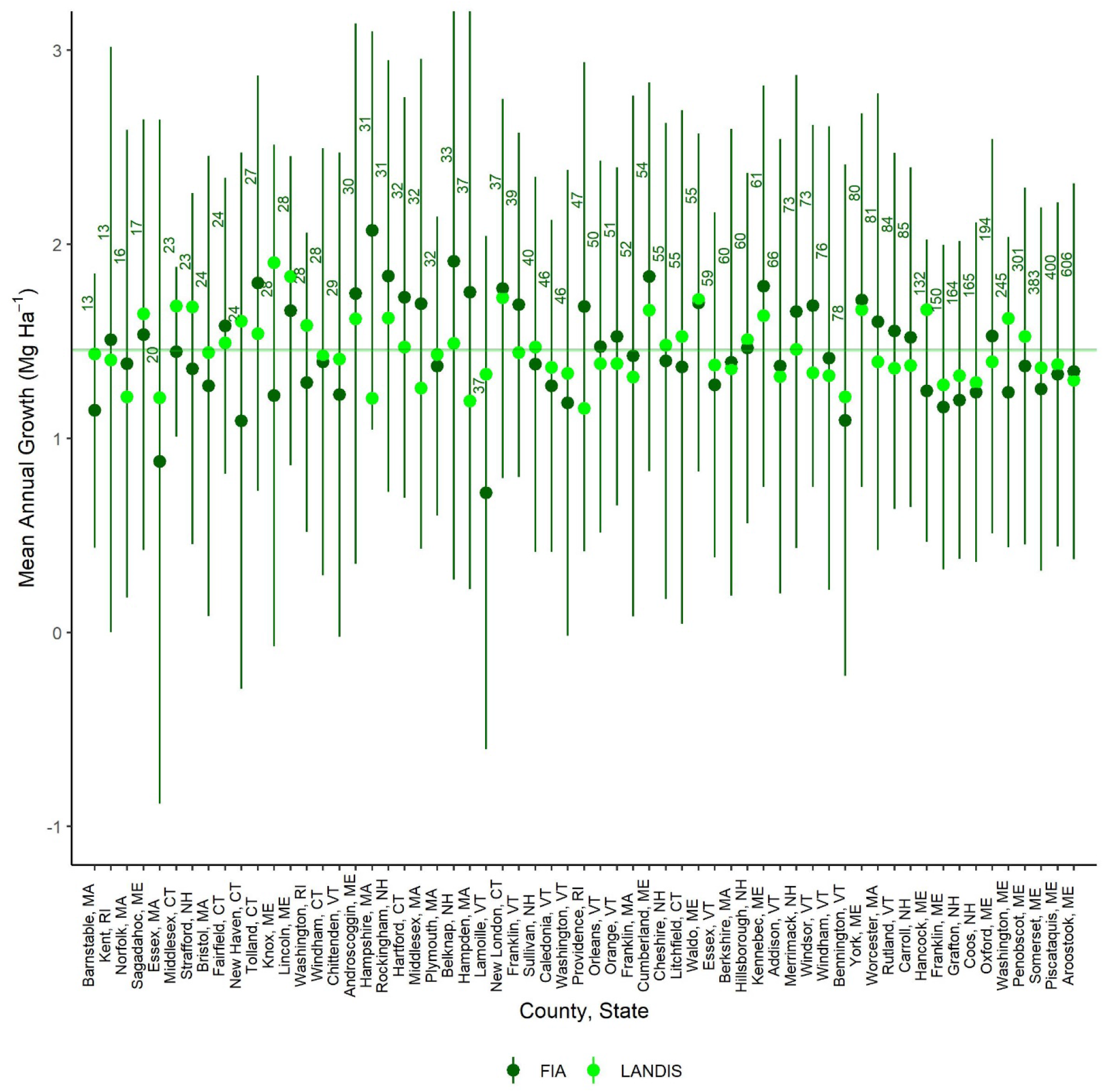
Observed carbon growth (dark green; FIA) and simulated carbon growth (light green; LANDIS-II) within New England counties with greater than 10 FIA plots. Dots and lines represent means and standard deviation, respectively, for the FIA data. Horizontal lines represent the grand means of both observed and simulated growth across counties, however they are insignificantly different (*p* < 0.05) and too close to be distinguishable.

#### HARVESTING

We used the LANDIS-II Biomass Harvest extension (v. 4.2) (Gustafson et al. 2000) at 10-year time steps to simulate timber harvest. We leveraged previous work by Duveneck and Thompson (2019) to define our harvesting prescriptions and initialize our allocation of those prescriptions for the Recent Trends (RT) scenario (Appendix II). For each alternative scenario, we adjusted the RT harvesting prescriptions and rates based on the stakeholder designed NELF scenario storylines (see below and Appendix II for specifics).

Several improvements to our modeling framework resulted in differences between our previous simulations of recent trends (Duveneck and Thompson 2019) and those presented here. Improvements include an updated version of PnET-Succession that does not initialize cohorts by growing each individual species-cohort. Rather, we used a recently developed function that gave each cohort a predetermined initial biomass based on the imputation of FIA plots into individual forest cells (from Duveneck et al. 2015). Specifying the initial biomass of each species-cohort reduced the uncertainty of our starting conditions and provided a consistent and better approximation of forest conditions at the beginning of each simulation. While updating our initial conditions to include initial biomass, we also simplified our initial communities and updated species-specific parameters. Compared to the results presented in Duveneck and Thompson (2019), these updates resulted in 9% more overall biomass in 2060 and only slight differences in relative species abundances.

We also improved our approach to simulating regional variation in management and the impacts of conservation on spatial harvesting patterns. To simulate regionally-specific harvest behaviors, we delineated ‘Management Areas’ as specific ownership groups and conservation statuses within New England states (Duveneck and Thompson 2019). Initially management regions were designated at the state level, but due to significant differences in both current harvest characteristics and changes described in the NELF scenario narratives, we split New Hampshire and Vermont into north and south regions to allow sub-state regional variation in harvest rates (see Appendix III).

To incorporate conservation in our modeling of harvest, we prohibited harvest in areas designated as conserved with USGS Gap Analysis Program (GAP) Status Codes 1 and 2, which represent conserved lands with management restricted to conservation purposes only (i.e., no commercial harvesting). We allowed harvest to occur on all other conserved lands, which is consistent with most multiple-use conservation restrictions. As areas changed within each scenario simulation from not conserved/restricted to conserved with GAP Status Codes 1 & 2, harvesting was reallocated from these newly conserved areas to forests that were not conserved with harvesting restrictions. We did this by defining a new set of management areas based on management region (i.e., state or substate area) and time step of conversion to conserved forest. During the time steps prior to conservation, the harvest rates and allocations for the conserved forest management areas were the same as those in the unconserved forests in that management region; then, at the time step of conservation, harvest rates were set to zero for the conserved forest management area and the rates of harvest were proportionally increased, based on area, for the unconserved parts of the management region (outside of the conserved forest management area). In this way, target harvesting rates were still met for each timestep of the simulation, but harvesting did not occur within areas projected to be conserved with GAP Status Codes 1 & 2. Thus, the effects of conservation did not have large effects on harvest rates at the landscape scale, as those rates remained true to the scenario storylines, but the spatial allocation of those harvests did change.

### Allocating harvest prescriptions for recent trends

To estimate the area to harvest in each management area, we used remeasured FIA plot data grouped by region and ownership type. Similar to the methods we used to parameterize forest growth and those in Thompson et al. (2017a), we used FIA plots with two or more measurements after 2000 to calculate the proportion of FIA plots harvested in each management area. The proportion of plots harvested of all available plots in a management area was then divided by the remeasurement period to estimate the annual harvest rate for each management area (See Appendix III). A plot was considered “harvested” if at least one tree was marked as removed within the FIA tree-level database between remeasurement periods. Therefore, we considered harvest in the broadest sense, including both commercial and incidental harvest (*sec*. Belair and Ducey 2018) in this analysis of harvesting. Similarly, to estimate average harvest intensity (i.e., percent biomass removed in a harvest), we joined FIA plot and individual tree data to calculate total carbon (C) for each plot and total and percent C removed through harvest between remeasurement periods. We then averaged the percent C removed in each management area to calculate the target average intensity of harvest for applying harvest prescriptions (Appendix III). Average harvest intensities were relatively low, since all types of tree removal were considered “harvests” for this analysis.

Within each management area, harvest prescriptions were implemented based on modified RT harvesting prescriptions from Duveneck & Thompson (2019) (Appendix II) and harvest proportions in Belair and Ducey (2018) (Appendix III). A single time-step test simulation of our model with the defined harvest prescriptions allowed us to compute the average harvest intensity (i.e., percent carbon removed) for each of the prescriptions. For these RT prescriptions, we then used a linear programming with maximum likelihood estimation method to determine the best allocation of harvest prescriptions within each management area so that the overall intensity of harvest in our simulations approximated the average harvest intensity from FIA for that management area (See Appendix III for more details).

### Carbon allocation

The fate of carbon removed from the landscape through harvesting was tracked using a common method for carbon accounting that was developed by the U.S. Forest Service for greenhouse gas accounting (Smith et al. 2006). We then adapted these carbon accounting methods to fit with our integrated modeling of aboveground carbon dynamics. While the Smith et al. (2006) carbon accounting methods were based on relatively older timber product output reports and mill efficiencies etc., the methods were both standard and flexible enough that we were able to modify these methods to use with the cohort modeling approach of LANDIS-II and PnET-Succession.

The Smith et al. (2006) carbon accounting methods track carbon from growing stock trees into several carbon pools (e.g., slash, landfill, firewood, and wood products) according to forest type and species-specific decay or transfer rates (Figure 5). These methods use individual tree measures (e.g., diameter, merchantability) to define growing stock, measures that are not simulated in LANDIS-II and PnET-Succession. Therefore, we modified the approach to accommodate the tree cohort outputs from LANDIS-II and cohorts 20 years old or older were considered potential growing stock. We used the Biomass Community Output extension in LANDIS-II (Scheller 2020) to evaluate cohort ages at the time of removal. For removed cohorts less than 20 years old (i.e. not potential growing stock and not tracked in the Smith et al. (2006) methods), 14% of the total carbon was allocated to the slash pool to account for material left on site to decay (following Reinmann et al. 2016), and the remaining 86% of the harvested carbon was allocated to the fuelwood category and was mineralized (emitted) by the next time step (Figure 5). Then, for all removed cohorts over 20 years old (potential growing stock), the same 14% was allocated to the slash pool to account for material left on site to decay, including trees that were not merchantable, with the remaining 86% of the removed cohorts considered ‘growing stock’, as used in Smith et al. (2006). The removed growing stock’s C was then allocated to different carbon pools at each time step using the modified Smith et al., (2006) accounting methods (illustrated in Figure 5, and in more detail in Appendix IV), with transfer and decay rates based on the forest type and wood type of the removed cohorts (Appendix IV). The harvested carbon allocation to different pools and decomposition rates were unaltered from the Smith et al., (2006) accounting methods for our RT scenario.

**Figure 5.**
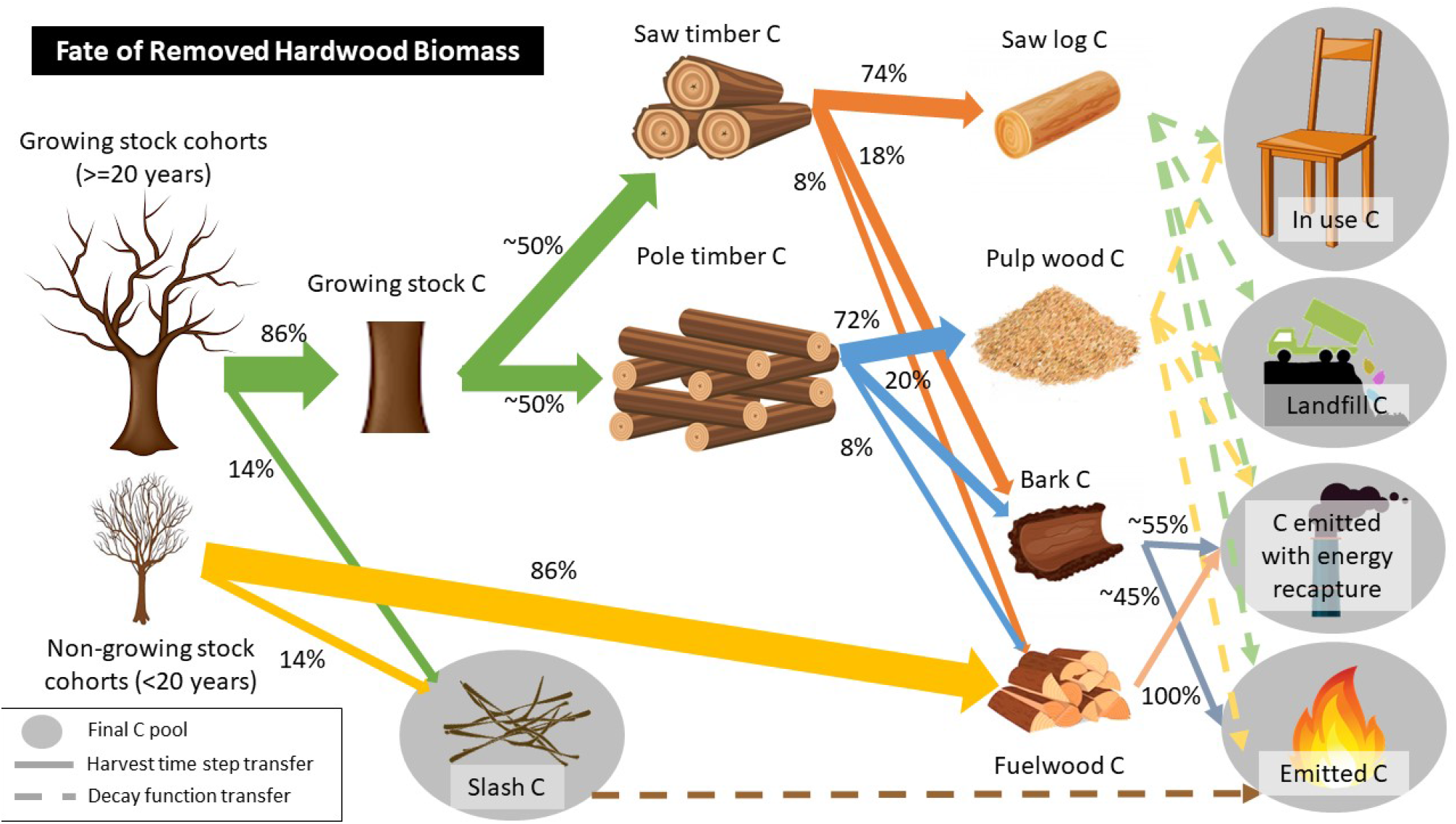
Example allocation of carbon into final carbon pools for hardwood species in the RT scenario. Proportions change for softwood species and by scenario. Dashed lines represent between-pool transitions, with allocation proportions dependent on time since removal, whereas solid arrows indicate transitions that are constant and occur at the time of removal.

Following a similar analysis by Reinmann et al., (2016), the carbon removed from development in RT was assumed to not enter the timber market. Instead, half of the carbon removed through development was allocated to fuelwood and mineralized (emitted) in that time step, and the other half of the removed carbon was added to the slash pool and was emitted using a softwood/hardwood specific decomposition rate (Russell et al. 2014). Note, our accounting framework only tracked carbon from harvesting or development during our simulation time-frame, from 2010-2060, so any carbon removed prior to 2010, or any transitions (e.g., from “in-use” to “emitted”) that happened after 2060, were not tracked.

### Translation of the scenarios into harvesting prescriptions and carbon allocation

Using the same methods as those used to translate qualitative stakeholder scenario descriptions of land cover change into quantitative inputs for our land-cover change model (Thompson et al. 2020), we translated the four NELF scenario narratives from qualitative descriptions of resource use and harvest patterns into differential rates of harvest intensity, area harvested, and carbon allocation (Appendices II and III). Each of the alternative scenarios had additional harvest prescriptions that were defined and directly linked to the scenario narratives and changes to harvesting rates were defined relative to Recent Trends (RT) (Appendix II). Some of the scenario narratives also indicated innovative approaches to development/timber use or energy generation, resulting in differential allocation of carbon into either in-use pools or emitted with energy recapture. For example, in Connected Communities (CC), stakeholders indicated a need to use biomass energy as a transition fuel to more renewable sources; this statement translated to the creation of a biomass harvest prescription where all biomass (minus that allocated to slash) was emitted with energy recapture.

## RESULTS

### Combined carbon consequences of land-use changes

Despite widely divergent land-use regimes, New England’s forests remained a net carbon sink to 2060—i.e., more carbon was sequestered in forests and stored in wood products than was released to the atmosphere—in four of the five future scenarios, including Recent Trends (RT) (Table 3, Figure 6). Only in the Go it Alone (GA) scenario did New England’s forests become a net carbon source, with total emissions of 68 Tg C, by the year 2060. Additionally, the amount of carbon stored in live biomass (i.e., sequestered) through 2060, was greater than the emissions from forestry and development in three of the five scenarios: RT, Connected Communities (CC), and Yankee Cosmopolitan (YC) (Figure 6). Only after accounting for the carbon stored in wood products, landfill, and slash did the Growing Global (GG) scenario become a net carbon sink over the 50 years, since carbon emissions in this scenario were greater than the carbon sequestered. In YC and CC, the lower amount of harvested carbon resulted in increased sequestration rates and reduced emissions as compared to RT. Increased harvesting in GA and GG resulted in nearly equal amounts of carbon stored and emitted. Below we describe in more detail the differences of contributions to each of the storage and emissions pools: live, stored, and development and forestry emissions.

**Table 3.** Total carbon emissions and storage for each scenario (storage includes the sequestered live aboveground forest carbon and any harvested carbon stored in wood, slash and landfills in 2060).

| Scenario | Total emitted (Tg C) | Total stored (Tg C) | Total carbon balance (Tg C) |
| --- | --- | --- | --- |
| Recent Trends (RT) | 360 | -672 | -312 |
| Connected Communities (CC) | 227 | -787 | -560 |
| Go it Alone (GA) | 574 | -506 | 68 |
| Growing Global (GG) | 482 | -526 | -44 |
| Yankee Cosmopolitan (YC) | 112 | -844 | -732 |

**Figure 6.**
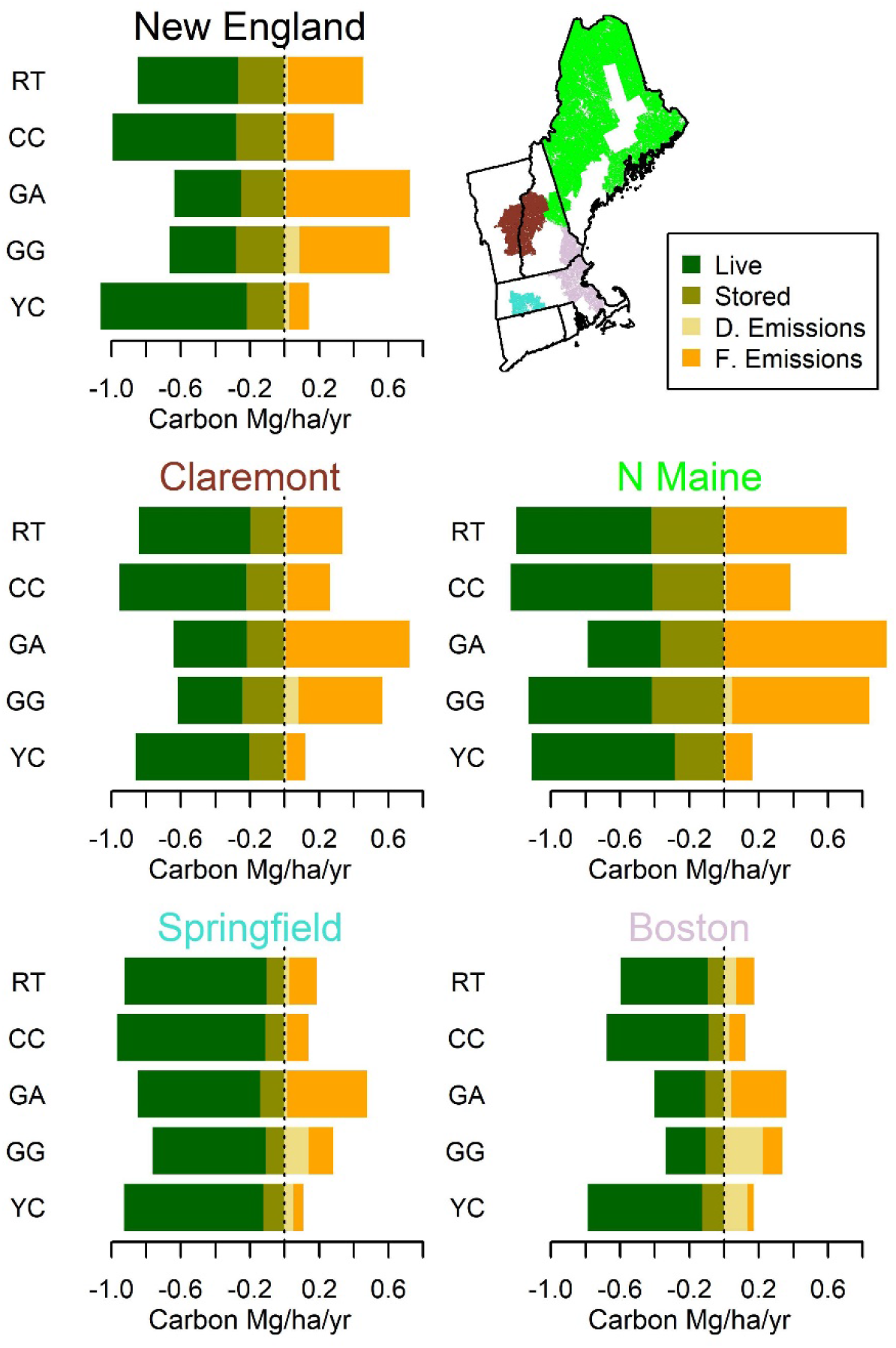
Rates of emission and storage for removed carbon and live carbon for all of New England, and within four example CBSAs: Claremont (in NH and VT), N Maine, Springfield in central MA, and Boston (which covers the seacoast in most of MA and NH). The colors of each CBSA name above each chart correspond to CBSA areas on inset map. “Live” represents the total carbon sequestration or accumulation of live biomass; “Stored” is the rate of storage of carbon in slash, wood products, and landfills; “D. Emissions” are the development emissions; and “F. Emissions” are the emissions from forestry for the full 50-year simulation.

### Forest carbon stocks

Forest growth in New England was the primary contributor to carbon storage in all scenarios, though there were regional/CBSA variations by scenario (Figures 6 & 7). These regional differences in live carbon stocks were not only driven by changes in land-use drivers, but also by climate, with warming enhancing growth more in the south than the north (Figures 6 & 7). In both CC and YC, forests accumulated more aboveground carbon (AGC) than in RT, generally from a combination of reduced timber harvesting and forest conversion (Figure 7). However, the increased harvesting and development reduced the ability of the forest to store carbon in both the GG and GA scenarios (Figure 7). The narratives of each of the scenarios also altered the spatial allocation of land use and therefore carbon. In the two global socio-economic connectedness scenarios, YC and GG, the impacts of harvesting and conversion are very similar, yielding higher losses of aboveground carbon nearer to currently highly developed areas (e.g. Boston, MA) and therefore less carbon accumulation/sequestration (Figure 7). Conversely, timber harvesting was a main driver of aboveground carbon removal in CC and GA, which resulted in less AGC accumulation in the less densely developed parts of New England (e.g., N Maine).

**Figure 7.**
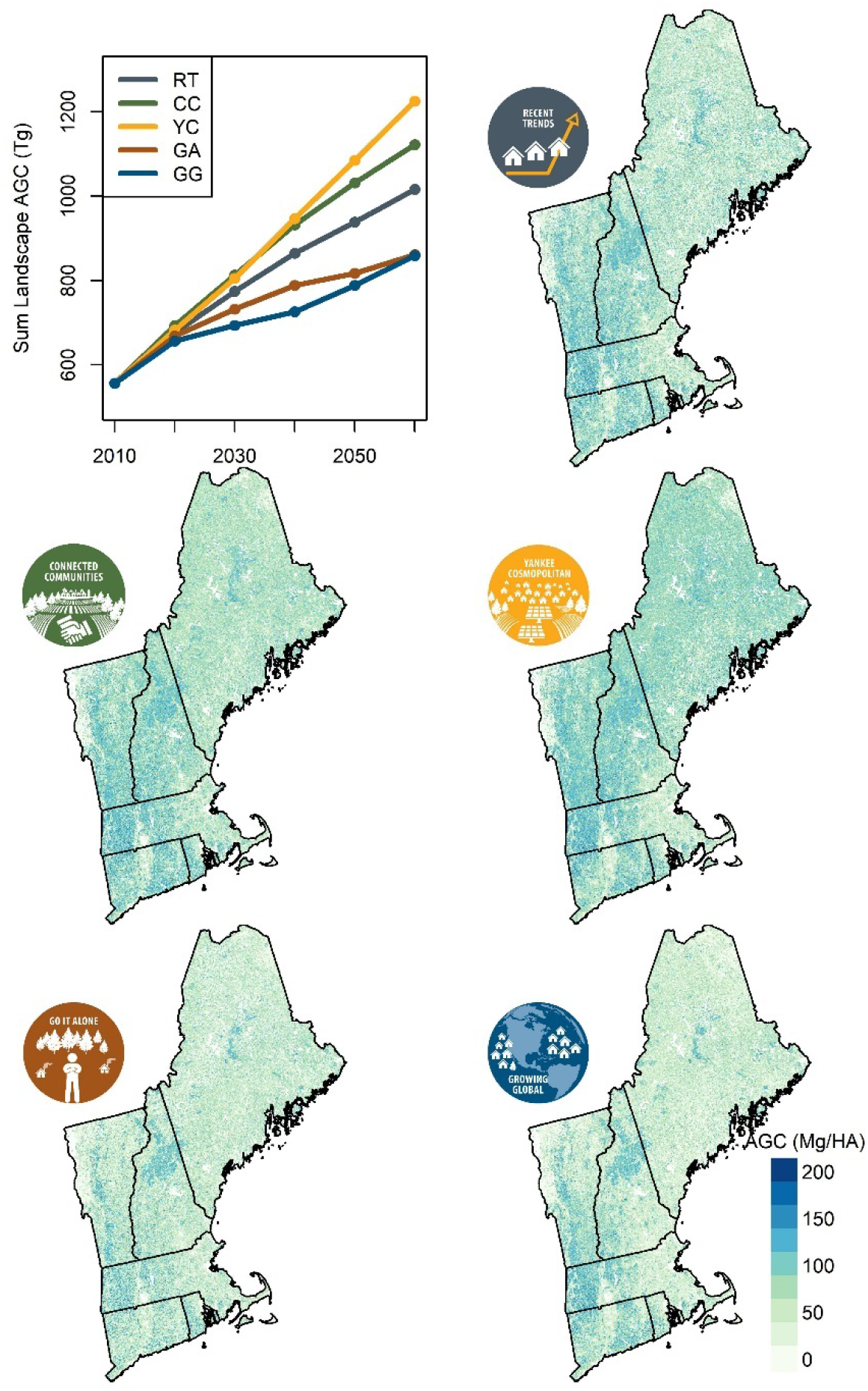
Maps of AGC (Mg ha^−1^) for each scenario at 2060. For comparison, Figure 3 shows AGC at year 2010 starting conditions. Line graph shows sum of AGC (Tg) accumulation for each scenario over time.

### Harvesting and development rates

Carbon emissions and storage varied spatially based upon the differences in development and harvesting for each of the scenarios by region/CBSA (Figure 6 & 8). For example, Boston had relatively higher development emissions in scenarios with global socio-economic connectedness (i.e., YC and GG) (Figure 6). In contrast, emissions from harvesting were higher in rural regions like Northern Maine for scenarios with local socio-economic connectedness (i.e., GA and CC) (Figure 6). Similar to previous studies (e.g., Canham et al. 2013, Thompson et al. 2017a, Duveneck and Thompson 2019), more C is removed through timber harvesting than through conversion of forests to development in all of the scenarios. Indeed, in the RT scenario presented here, 12x more carbon was removed by harvesting than by development (Figure 8). Importantly, three of the four stakeholder-articulated scenarios predicted an increase in harvested area, but the intensity and spatial allocation of harvesting were distinct in each scenario (Appendix III).

**Figure 8.**
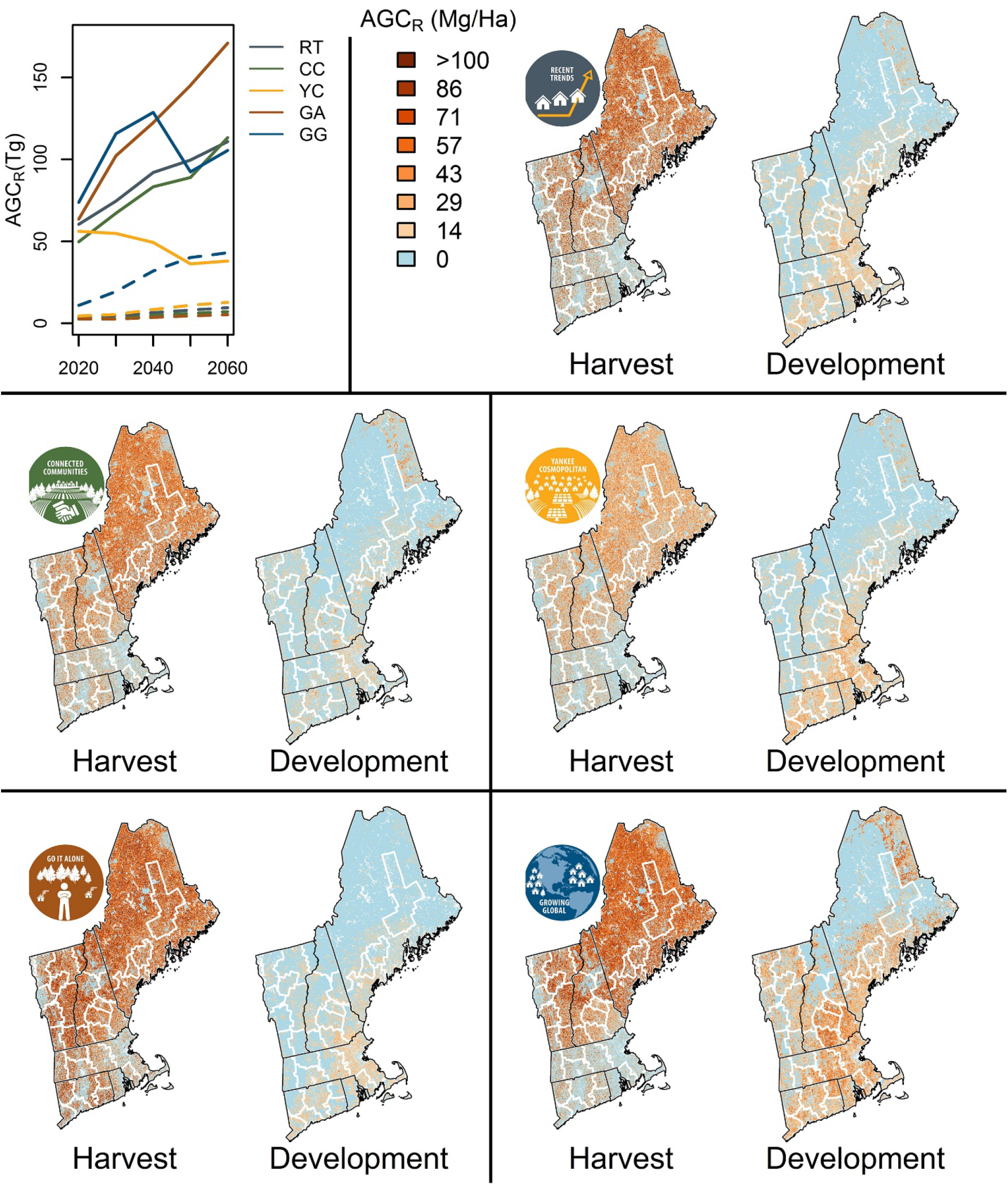
Line chart shows total aboveground carbon removed (AGC_R_) over time for each scenario. Harvest removals are solid lines. Developed removals are dashed lines. Maps of total removed aboveground carbon by either harvest or development for each scenario with CBSAs outlined in white and state boundaries outlined in black.

Given the increase in the target harvested area outlined in all but the YC scenario, some of the management areas did not have enough forested area that met harvest criteria remaining in 2060 to sustain harvest rates. Therefore, some scenarios deviated in total area harvested from the harvest area targets. Specifically, as shown in Figure 8, the GG scenario did not have enough suitable stands available to meet the target harvest area beginning in 2040. However, although the GA scenario had similar harvest area targets, our models were able to continue to harvest at nearly the target rates throughout the simulation by allowing more harvest to occur in southern New England, whereas GG limited harvesting to the northern reaches of NE (Figure 8). The resulting total harvested area after 50 years for GG was 143% of RT and the area harvested in GA was 144% of RT. Similarly, CC harvested 129% of the total area harvested in RT. Only the YC scenario resulted in less area harvested, approximately 79% of the area harvested in the 50-year RT simulation (Figure 9a).

**Figure 9.**
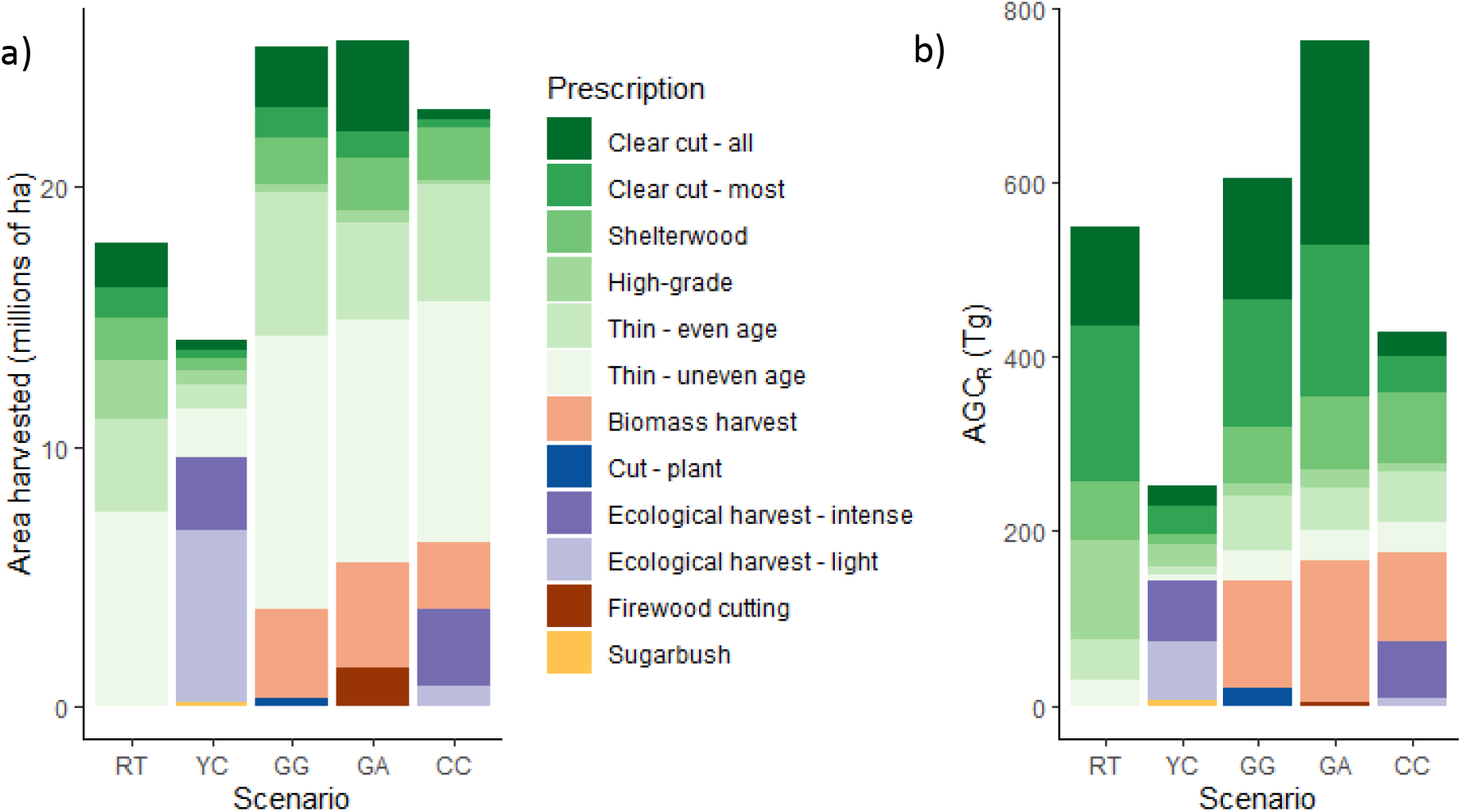
a) Cumulative area harvested by prescription for each scenario. b) Cumulative aboveground carbon removed by harvest prescription for each scenario. Prescriptions shown in green are the original Recent Trend prescriptions, while the other prescriptions are those created and defined from the alternative scenario narratives.

Total carbon removed by harvest varied by scenario and the intensity of the alternative harvest prescriptions defined in the scenario narratives. New scenario-specific prescriptions (i.e., not used in RT) were generally less intense than those in RT (Table 4) and often emulated attributes of silvicultural practices that promote diversity and potentially longer-term carbon storage (e.g., longer rotation periods, promoting/retaining a diversity of age, size, and species). As a result of these new prescriptions, both of the high natural resource planning and innovation scenarios, CC and YC, removed less overall carbon from the landscape than RT (CC removed 78% of RT, and YC removed 46% of RT), despite CC harvesting more area (Figure 9). GG and GA both removed more C in the form of harvested timber than RT (110% and 139% of RT, respectively), and the difference between these two scenarios was primarily driven by differences in the intensities of the applied harvest prescriptions (Figure 9, Table 4).

**Table 4.** Changes in area and intensity of development and harvesting by scenario. Harvest intensity includes all types of tree removal – commercial and incidental (non-commercial). Development intensity reflects an assumption that forested sites converted to agriculture, high-density development, and low-density development will reduce forest biomass by 100%, 94%, and 50%, respectively.

| Scenario | Total Area Developed/<br>Converted<br>(K ha)<sup>1</sup> | Average Development<br>Intensity (%<br>AGC<sub>R</sub>) | Total Harvest<br>Area (K ha) | Average Harvest<br>Intensity<br>(% AGC<sub>R</sub>) |
| --- | --- | --- | --- | --- |
| Recent Trends<br>(RT) | 567 | 60% | 17853 | 35% |
| Connected<br>Communities<br>(CC) | 325 | 82% | 23036 | 25% |
| Go it Alone (GA) | 375 | 53% | 25645 | 36% |
| Growing Global<br>(GG) | 2199 | 71% | 25450 | 32% |
| Yankee<br>Cosmopolitan<br>(YC) | 170 | 58% | 14119 | 25% |

### Harvesting and development emissions and storage

In the RT scenario, approximately two-thirds of the removed carbon, from either harvesting or development, was emitted by 2060, totaling 360 Tg C (Figure 10). One-third, or 212 Tg C was stored in use, landfilled, or as slash. The fate of removed carbon for the alternative NELF scenarios differed based on how the narratives described carbon emissions and storage deviated from RT. For example, in scenarios with high natural resource planning and innovation (i.e., CC and YC), the narratives described increased use of wood products and decreased landfilling of wood products, keeping a higher proportion of the removed carbon in storage by the year 2060 than in other scenarios (Figure 10). These scenarios also had fewer total carbon emissions than RT and a more balanced allocation of carbon into emitted vs. stored pools, with YC having the lowest overall emissions at 112 Tg C and approximately 61% (174 Tg C) of the removed carbon remaining in stored pools at the end of the simulation. CC had nearly equal proportions of carbon emitted and stored in 2060, with emissions of 227 Tg C and 221 Tg C stored. Both of the scenarios with lower natural resource planning and innovation had much higher emissions, both as a proportion of total carbon allocation and total carbon emitted. Sixty-eight percent of the carbon removed in GG was emitted by 2060, totaling 482 Tg C (with 223 Tg C stored), and 74% of the carbon removed in GA was emitted, totaling 574 Tg C (with 201 Tg C stored) (Figure 10).

**Figure 10.**
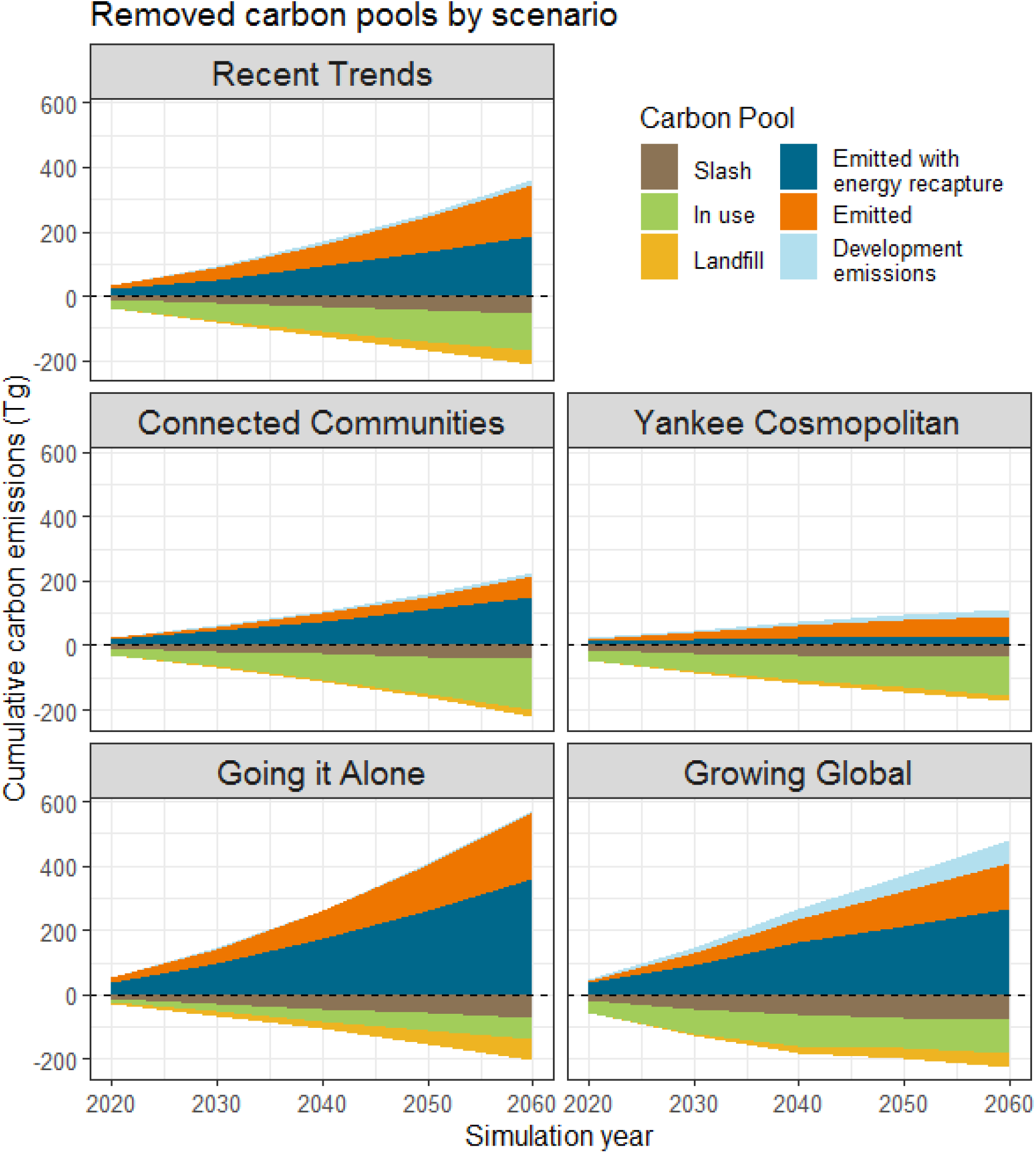
Cumulative total carbon emissions and storage from removed carbon for each scenario throughout the simulation. Additional breakdown of emissions by removal type is in Appendix IV.

## DISCUSSION

In forests around the world, land-use decisions will influence whether a landscape will act as a net carbon sink or source. Indeed, the land-use decisions in each of our scenario narratives drove whether New England forests remained a net carbon sink or source in our 50-year simulations. In some scenarios, like Yankee Cosmopolitan (YC), the recovery dynamics of the relatively young New England forests and increased growth due to climate change allowed forests to remain a strong carbon sink. However, in others, such as Go it Alone (GA), the individual management choices of private forest landowners produced carbon emissions that surpassed the ability of New England forests to sequester carbon, and New England forests became a net carbon source by 2060. The impact of the individual scenarios on carbon dynamics was most closely tied to changes in “natural resource planning and innovation” within each of the narratives. Along this axis of change, stakeholders described changes to harvest intensity, area harvested, as well as how much of the harvested timber went into long-term storage as compared to the Recent Trends (RT) scenario. These changes in timber production and use substantially altered the carbon balance of New England in 2060.

In the RT scenario, there was an additional 312 Tg C stored in 2060, as compared to 2010, the start of the scenarios, primarily stored as live biomass within forests. Forest growth in the RT scenario, enhanced by climate change, resulted in an increase in carbon stocks by 670 Tg C as compared to starting conditions. Of the removed carbon in RT, approximately two-thirds of it was emitted into the atmosphere by 2060, with 95% of these emissions from harvesting (both commercial and incidental). These emissions are equivalent to 1,320 MMtCO_2_eq over the 50-year simulation. Our estimates of carbon impacts of land use in the RT simulation are comparable to other studies of carbon change. For example, Harris et al. (2016) estimated that New England as a whole stored 16.1 Tg C yr^−1^, emitted 9.0 Tg C yr^−1^; for a net carbon change of - 7.1 Tg C yr^−1^ (negative indicating stored) between 2006 and 2010. In our projection of recent trends, New England stored around 13.4 Tg C yr^−1^ and emitted 7.2 Tg C yr^−1^, for a net carbon change of −6.2 Tg C yr^−1^. Like in Harris et al. (2016), the vast majority of our projected carbon losses (i.e., emissions) were from harvesting, but the continuation of New England forests’ recovery from mid 19^th^ century deforestation and increased growth due to climate change (Duveneck et al. 2017) resulted in a net increase in carbon stocks. Therefore, projected changes in harvesting for each of the stakeholder defined NELF scenarios had the largest impacts on carbon stocks and fluxes through 2060.

When designing the scenarios, the stakeholders tried to envision changes to harvesting practices and wood product utilization that diverged quite a bit from those in RT and each other. For example, in the Connected Communities (CC) scenario, stakeholders created a narrative that described a transition to ‘ecological forestry’, but an increase in overall harvested area. Therefore, despite harvesting nearly 30% more area compared to RT, the transition to ‘ecological forestry’ resulted in a 10% reduction in average harvest intensity, and therefore 125 Tg C less removed from the landscape. The narrative of CC also focused on innovative uses and valuing of local timber products as part of “natural resource innovation.” These new local timber products assumed new technologies like cross-laminated timber products for building materials (New England Forestry Foundation 2017, Kaboli et al. 2020), resulting in more of the removed carbon remaining in “in use” products by the end of the simulation. For this scenario (i.e., CC), the combination of the increase in timber that remained in durable goods and the reduction in overall harvest intensity resulted in 133 Tg less carbon emitted than in RT and 115 Tg more carbon stored in the same time period. The combined carbon benefit of these choices resulted in approximately 909 MMtCO_2_eq fewer emissions through reduced direct emissions and increased sequestration. Importantly, of the carbon emitted, there was only 34 Tg C less emitted with energy recapture than in RT, indicating that CC could continue to meet most of the projected wood energy demands of New England as in RT. Similarly, 43 Tg more stored carbon remained in use at the end of the simulation, meaning projected wood product demand could also be met at similar levels as RT.

Correspondingly, an overall reduction in harvesting, in both area and intensity, has an even more pronounced impact on carbon emissions and storage, as seen in the Yankee Cosmopolitan (YC) scenario. Since the YC narrative emphasized global connectedness, fewer natural resources needed to be sourced in New England than in CC or RT, allowing total harvesting to reduce dramatically in this scenario. Along with the reduction in timber harvesting, the YC scenario described landfilling less long-term wood products, which resulted in more carbon remaining in storage throughout the simulation. These land-use choices (i.e., YC scenario) resulted in the largest decrease in emissions, 248 Tg C less than RT, and the largest increased in C stocks, 172 Tg C more than in RT, primarily through increased forest growth. This had a combined carbon benefit of approximately 1,540 MMtCO_2_eq fewer emissions (and increased sequestration) as compared to RT.

However, land-use decisions such as those described in these NELF scenarios also have carbon consequences which were not represented in our simulations (e.g., issues of substitution or leakage). The carbon impacts of sourcing products, such as building materials, and changes to energy demand/production to meet the increasing population in YC were outside the scope of this analysis and yet have major carbon emissions implications. For example, in the YC scenario an additional 3 Tg C is “in use” at the end of the scenario, as compared to RT, but the housing demand is likely to be much higher in YC. Therefore, it is likely that these building materials would need to be sourced from other parts of the world, causing leakage not addressed in this paper (Henders and Ostwald 2012). Additionally, nearly 165 Tg C less was emitted with energy recapture in YC, meaning without meaningful energy efficiency measures, energy would need to be produced through other means, such as renewable sources (as described in the YC narrative), that would also have land use and carbon implications.

Conversely, stakeholders also described two scenarios that resulted in higher carbon emissions from land use than RT (GG and GA), and one where New England Forests became a net carbon source by 2060 (i.e., GA). The Go it Alone (GA) narrative described a future land-use scenario where New Englanders met local demand for wood products through increased local harvest, increasing total area and harvest intensity, and relied more heavily on biomass energy (as opposed to acquiring electricity or heat from distant power-plants). These two changes to land use and energy generation resulted in a scenario that emitted 68 Tg more carbon than it sequestered and stored over the 50-year simulation. As compared to RT, GA emitted 214 Tg more carbon, especially in the emissions with energy recapture pool (e.g., biomass energy), and stored 166 Tg C less, with a combined net increase in emissions of approximately 1,393 MMtCO_2_eq. While these land-use choices resulted in a scenario where forests were unable to sequester carbon at a rate greater than the emissions from harvesting, the 358 Tg C emitted with energy recapture in GA may offset some emissions from other energy sources, such as fossil fuels, though the benefit from replacement of these fuel types was outside the scope of this study.

Similarly, Growing Global (GG) also expanded total harvest area to meet higher demand for wood products due to a quickly increasing human population (as described in the GG narrative). The combination of the increase in harvesting and development in GG resulted in 122 Tg C more emissions than RT and 146 Tg C less storage, contributing a net increase in emissions of 983 MMtCO_2_eq over the 50-year simulation, as compared to RT. Despite GG having the largest expansion of developed area of any scenario, increasing the total development 288% over RT, harvesting was still responsible for over 85% of the total carbon emissions. Despite the overwhelming contribution of harvesting to emissions, development negatively impacted sequestration. Indeed, simulated harvest generally resulted in slightly increased rates of sequestration in the 50 years of our study (though lowered stocks), while development resulted in both the reduction of stocks and no sequestration at that site. As visualized in Figure 6, development in GG caused rates of sequestration to be similar in GG and GA, despite significantly more tree removal in GA.

The GG scenario described a rapid expansion of total harvested area and a larger proportion of the simulated harvested timber remaining “in use”, or stored, as building materials due to the rapidly expanding development. However, we found that the forested area in GG was not able to sustain the high levels of harvesting needed to meet the increased demand in our 50-year simulations. These results extend what other recent studies have found, which is that current levels of timber harvesting are creating degraded and poorly stocked forests in New England, particularly in the northern-most areas where harvesting rates are highest (e.g., Gunn et al. 2019). Since most of the harvesting for GG was targeted for the more rural, northern areas of New England, the already degraded forests could not meet the demand for building lumber. Therefore, the simulated total harvested area was approximately the same as in GA, with slightly lower average intensity harvests. The timber harvesting described in the original GG scenario was therefore not sustainable, and also could lead to further carbon emissions due to the need to meet these demands using non-timber products or imported lumber.

Changes to timber harvesting and use, as well as development, had individual and interactive impacts on total carbon storage and emissions in New England. However, harvesting had the most immediate and profound effects on total emissions and the ability of the forests to sequester and store carbon. Interestingly, it was the combination of stakeholder described changes in both harvest area and intensity that drove changes to total carbon removed. The two extractive scenarios, GA and GG, described rapidly expanding harvest areas at current intensity levels and resulted in higher emissions and lower sequestration than RT. However, YC and CC described a decrease in overall harvest intensity, but CC was matched with an increase in total harvested area. These two scenarios (i.e., YC and CC) with less intense harvests sequestered more carbon than the other scenarios, including RT.

While overall harvesting drove most of the changes in simulated carbon sequestration and storage, the uses of the cut timber altered the proportion of the removed carbon remaining in stored pools at the end of each scenario. For example, in the RT scenario, by the end of the 50-year simulation, approximately 66% of the wood was emitted, but in the CC scenario, which focused on using wood in innovative long-term durable goods (e.g., cross-laminated timber), only approximately 50% of the harvested carbon was emitted by 2060. These scenarios show the importance of both decreasing harvest intensities and increasing long-term wood product storage as two measures for increasing carbon storage and sequestration and reducing land use carbon emissions. For the most immediate impacts on climate change and reduction of atmospheric CO_2_, land-use decisions that reduce total carbon removed from the landscape (e.g., reductions in harvesting) have the largest potential to reduce emissions and increase storage. However, for these short-term gains in forest carbon to be true gains, these land-use decisions must be paired with reductions in the consumption of wood products and their replacements that would otherwise lead to leakage and substitution emissions of equal or greater impact.

### Limitations and future directions

These scenarios and simulations present the carbon implications for land-use decisions that may occur by 2060. However, these decisions will impact carbon storage for years beyond the end of our simulations. For example, while the impacts of development on carbon for each of our scenarios was limited, the permanent conversion of land from forest to development has long-term impacts on sequestration that would not be borne out in the timeframe of these scenarios (Sleeter et al. 2018). We expect that over longer timeframes, the impact of development in these scenarios would become more pronounced. Additionally, we did not explicitly quantify the forgone sequestration from development, or the carbon accumulation that would have occurred if the development had not. Our simulations do quantify the direct impacts of harvesting and development on sequestration through their cumulative impacts on final carbon balance, but we did not quantify the indirect impacts of land use on the carbon potential of the landscape. We expect that including forgone sequestration would increase indirect carbon emissions from development, though the magnitude of this impact should be explored in further research. Similarly, our simulations do not account for emissions from sources that were created prior to 2010. For example, slash from harvests prior to 2010 were not a source of carbon emissions in our carbon accounting framework. Finally, we acknowledge that belowground carbon is an incredibly important aspect to carbon accounting, encompassing approximately half, or more, of the landscape carbon, with its own complex spatial patterns (Woodall et al. 2015, Jevon et al. 2019, Finzi et al. 2020). These spatial complexities can also emphasize the differential impacts of development and timber harvesting, but given this complexity and our modeling framework, we felt addressing the potential shifts in belowground carbon were beyond the scope of this paper.

Another limitation is that the carbon accounting framework used for the Recent Trends scenario is based on timber product reports, markets, and technologies that were available nationwide in the early 2000s (Smith et al. 2006). We expect that due to changes and improvements in timber production, these methods may now slightly underestimate the total amount of timber that is “in use” at the end of the simulation and overestimate the total emissions. While the magnitude of the effect of the timber production improvements is unknown, other carbon accounting methods give similar results (Harris et al. 2016), indicating that the overall effect on our carbon accounting is likely small and the relative changes of emissions and storage in the scenarios are still pertinent.

We also did not directly try to model changes to emissions and storage in each scenario using specific technology (e.g., housing changes, cross-laminated timber, smaller saw-kerf), since it is difficult to predict what technologies will be most relevant or may exist in 2060. Instead, we tried to account for these changes by implementing relative changes to what stayed in long-term wood products in our carbon accounting framework. In addition, we did not explicitly account for carbon leakage and substitution (i.e., the carbon emissions from products that would need to be garnered from new sources or locations given a reduction in the availability of timber), although these would impact overall carbon emissions for each of the scenarios. Finally, we did not address the myriad of other benefits forests have in the region, many of which have been explored in other papers using these scenarios (e.g., Thompson et al. 2020, Pearman-Gillman et al. 2020a, 2020b, Guswa et al. 2020), instead limiting our focus to the direct carbon impacts of land use. We hope that these scenarios will continue to be used to explore the impacts of future land-use decisions on other ecosystem services.

This work highlights how even seemingly small land-use decisions can have major impacts on the ability of the forests to mitigate climate change. For example, the 10% reduction in harvest intensity, coupled with the increase in long-term storage of wood products in the Connected Communities scenario resulted in emissions reductions that are equivalent to taking all of the passenger cars in New England off the road for nearly 30 years (FHWA 2015). Importantly, much of the reduction in harvest intensity in the Connected Communities scenario was implemented in northern New England. Here, parcels are larger, forest ownership is more focused on timber, and forests have more potential for additional carbon sequestration through enhanced silvicultural strategies (Thompson et al. 2017a, Gunn et al. 2019, Cook-Patton et al. 2020b). Additionally, by engaging in thoughtful regional planning to avoid rapid expansion of development like that simulated in Growing Global, we can also keep forests as forests and ensure these lands continue to sequester carbon into the future. As we work to promote resilient forests that can help mitigate the impacts of climate change, this research supports keeping as much of the land forested as possible, implementing sustainable harvest practices that maximize diversity and carbon storage through well planned management, and investing in technologies that encourage longer-term storage of wood products.

## ACKNOWLEDGEMENTS

We would sincerely like to thank Kathy Fallon Lambert who was integral in the design of the New England Landscape Futures project and scenarios. We would also like to thank Dr. Eric Gustafson for his insightful comments on a draft of this manuscript. This research was supported in part by National Science Foundation funded to the Harvard Forest Long Term Ecological Research Program (Grant No. NSF-DEB 12-37491) and the Scenarios Society and Solutions Research Coordination Network (Grant No. NSF-DEB-13-38809).

## APPENDICES

## APPENDIX I NELF Scenario Creation and Land Cover Change Simulation

The NELF scenarios were developed using the intuitive logics 2-by-2 matrix approach, popularized by Royal Dutch Shell/Global Business Network (Bradfield et al. 2005). In a series of six one-day workshops held throughout New England, stakeholders were guided through a structured process to identify and agree upon two drivers of landscape change that they deemed to be the most impactful and uncertain. The extreme conditions of these two drivers were then used to create a matrix with four quadrants that correspond to four scenarios. The two drivers used to create the NELF scenarios were: Socio-economic connectedness (local to global) and natural resource innovation (low to high). After identifying the dominant drivers, the stakeholders built-out the scenarios, incorporating their subsidiary drivers and initial descriptions of land use, into ~1000-word narrative storylines; attributes of each scenario are shown in Figure 1. Participants were then presented with a summary of recent land-use trends and asked to describe how land use would differ in each of the alternative scenarios using semiquantitative terms. In the months following the workshops, the NELF team reconvened the stakeholders in a series of interactive webinars to define the amount, intensity, and geography of land cover change in the scenarios.

The New England Landcover Futures (NELF) scenarios narratives were then translated into quantitative rates of land cover change and simulated using a spatially explicit cellular automata model called Dinamica Environment for Geoprocessing Objects (Dinamica – EGO). Initially, the NELF Recent Trends scenario was parameterized using historical rates and patterns of land cover change from 1990-2010. These parameters were derived via classified remotely-sensed Landsat imagery, specifically a timeseries of land cover maps created using the Continuous Change Detection and Classification (CCDC) algorithm (Zhu and Woodcock 2014, Olofsson et al. 2016). The four stakeholder scenarios: Connected Communities, Yankee Cosmopolitan, Go it Alone, and Growing Global were also simulated with Dinamica - EGO, and were based on modifications to the rates and spatial allocation of land cover transitions in the Recent Trends scenario. For more information on how the Dinamica - EGO model operates, see Thompson et al. 2017b. For more information how the NELF scenario narratives were translated into model inputs, see Thompson et al. 2020.

## APPENDIX II Defining harvest prescriptions

Harvest prescriptions and rates were initially based on the continuation of ‘recent trends’ in harvesting, following on the work done by Duveneck and Thompson (2019) alongside forestry professionals. Additional harvest prescriptions were defined based on the specific scenario narratives and current practices in forestry (Table A1 and A2). Scenario narratives were translated from stakeholder quotes to both new prescriptions, as well as changes in overall rates of harvesting and spatial allocation of harvesting (Table A2). Please see Appendix III for the rates and spatial allocation of harvests.

**Table A1.** Harvest prescription descriptions.

| Harvest prescription | Definition |
| --- | --- |
| **Clear cut - all** | All cohorts of all species are harvested, (the most intensive harvest prescription). The site must not have been cut in the last ten years and must have a cohort at least 50 years old to be eligible. |
| **Clear cut - most** | 80% of cohorts less than 20 years old remain in the harvest area, while all other cohorts are removed. The site must not have been cut in the last ten years and must have a cohort at least 50 years old to be eligible. |
| **Shelterwood** | Removing most of the cohorts (60% of all species >20 years of age) to regenerate species in partial shade. No species preference in the prescription. The site must not have been cut in the last ten years and must have a cohort at least 80 years old to be eligible. |
| **High-grade** | Only species of high value are 100% removed (see below for list) at varying ages depending on value, and all others are left. The site must not have been cut in the last ten years and have the most valuable species to be eligible. |
| **Thin – even age** | Higher intensity thinning that targets younger cohorts (<130 years old (y) removed at 30%), but all species equally. Older cohorts (>=130 y) are removed at lower intensities (5%). The site must not have been cut in the last ten years and have cohorts at least 40 years old to be eligible. |
| **Thin – uneven age** | Lower intensity thinning that would incorporate primarily non-commercial harvests and treats all species equally. Younger cohorts (<130 y) are removed at 7%, and older cohorts (>=130 y) at 5%. Any site with at least one 30-year-old cohort is eligible. |
| **Biomass harvest\*** | All species are removed at 50% or higher, with all less-valuable species removed. The site must not have been cut in the last ten years and have cohorts at least 60 years old to be eligible. All removed biomass (minus the slash component) is allocated to the energy emissions category during C accounting. |
| **Cut/plant\*** | Total clear cut and plant loblolly pine (*Pinus taeda*) as a crop tree species. The site must not have been cut in the last ten years and have cohorts at least 60 years old to be eligible. |
| **Ecological harvest – intense\*** | Aims to retain older cohorts, create longer rotation periods, create structural (age) diversity and regenerate species with higher economic and ecological value (e.g. oaks, pines, sugar maple, fir, etc.). The intense prescription takes out a higher % of biomass on average to continue to provide timber in a high demand market. |
| **Ecological harvest – light\*** | Aims to retain older cohorts, create longer rotation periods, create structural (age) diversity and regenerate species with higher economic and ecological value (e.g. oaks, pines, sugar maple, fir, etc.). The light prescription takes out a lower % biomass to retain more carbon on sight in scenarios where carbon is more important. |
| **Firewood cutting\*** | Very low intensity cutting (5%) of slightly older (>40 year old) primarily hardwood species. Any site with at least one 40 year old cohort is eligible. All removed biomass (minus the slash component) is allocated to the energy emissions category for C accounting. |
| **Sugarbush\*** | The site must be at least 75% sugar maple that is at least 50 years old (tap-able trees start at around 10" and sugarbush should have at least 74 taps/ha; Ferrell, 2013). Everything except for 20% of sugar maples 50-80 years, half of the sugar maples from 20-50, and 80% of red maples over 50 years is removed. The site must not have been cut in the last ten years and have cohorts at least 50 years old to be eligible. |
| * indicates a specialized prescription that was created based on at least one of the alternative land use scenarios. |  |

**Table A2.**
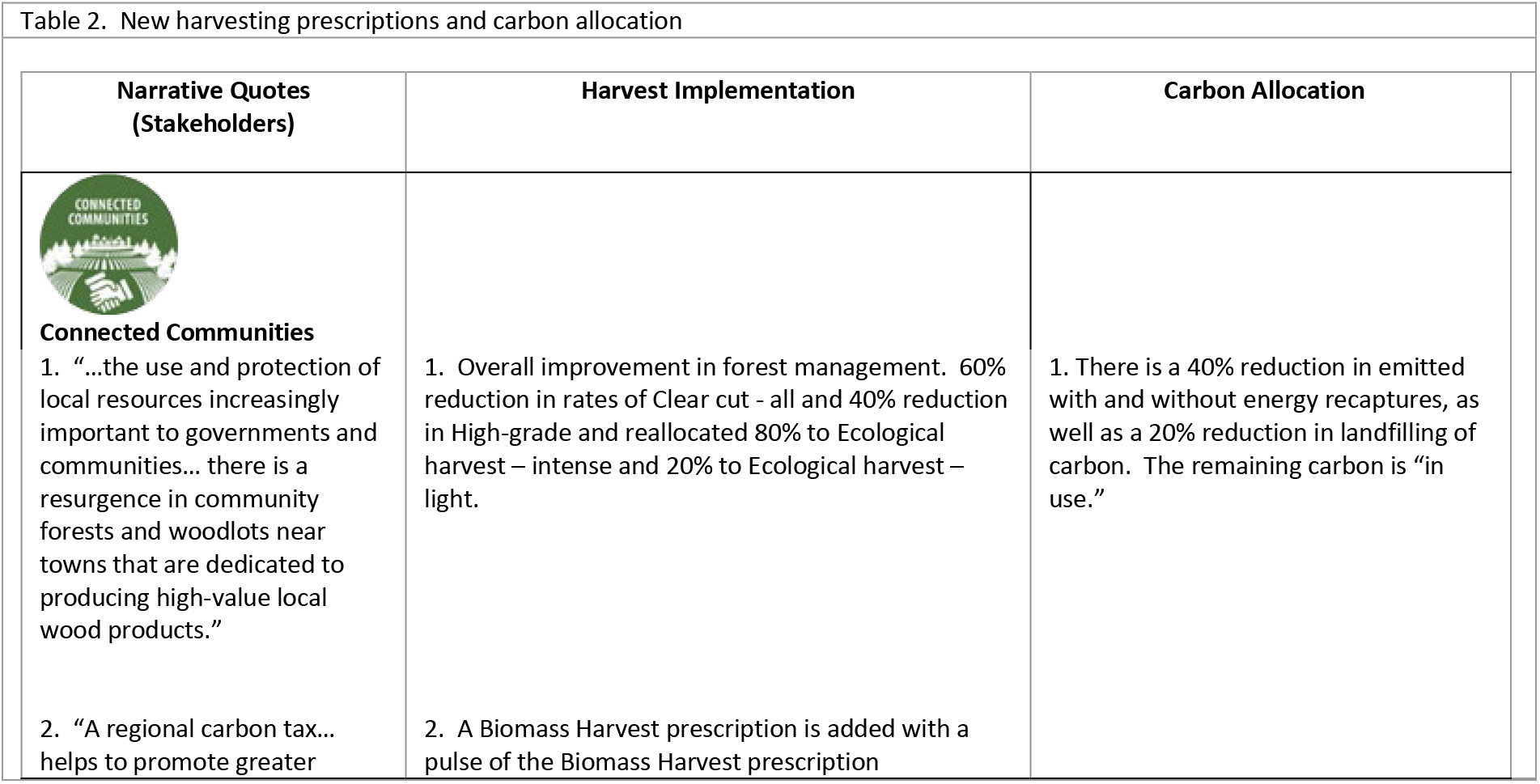
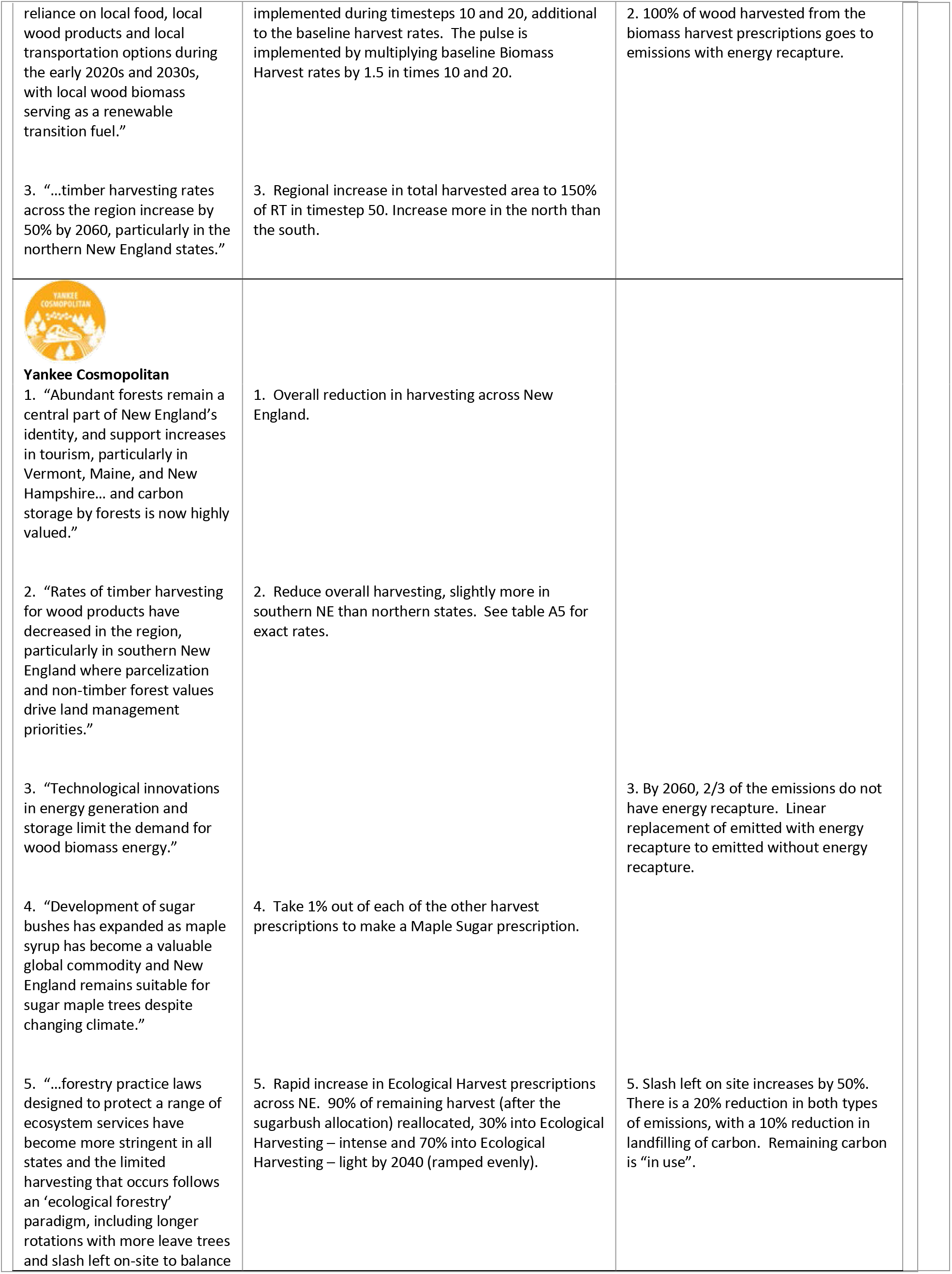
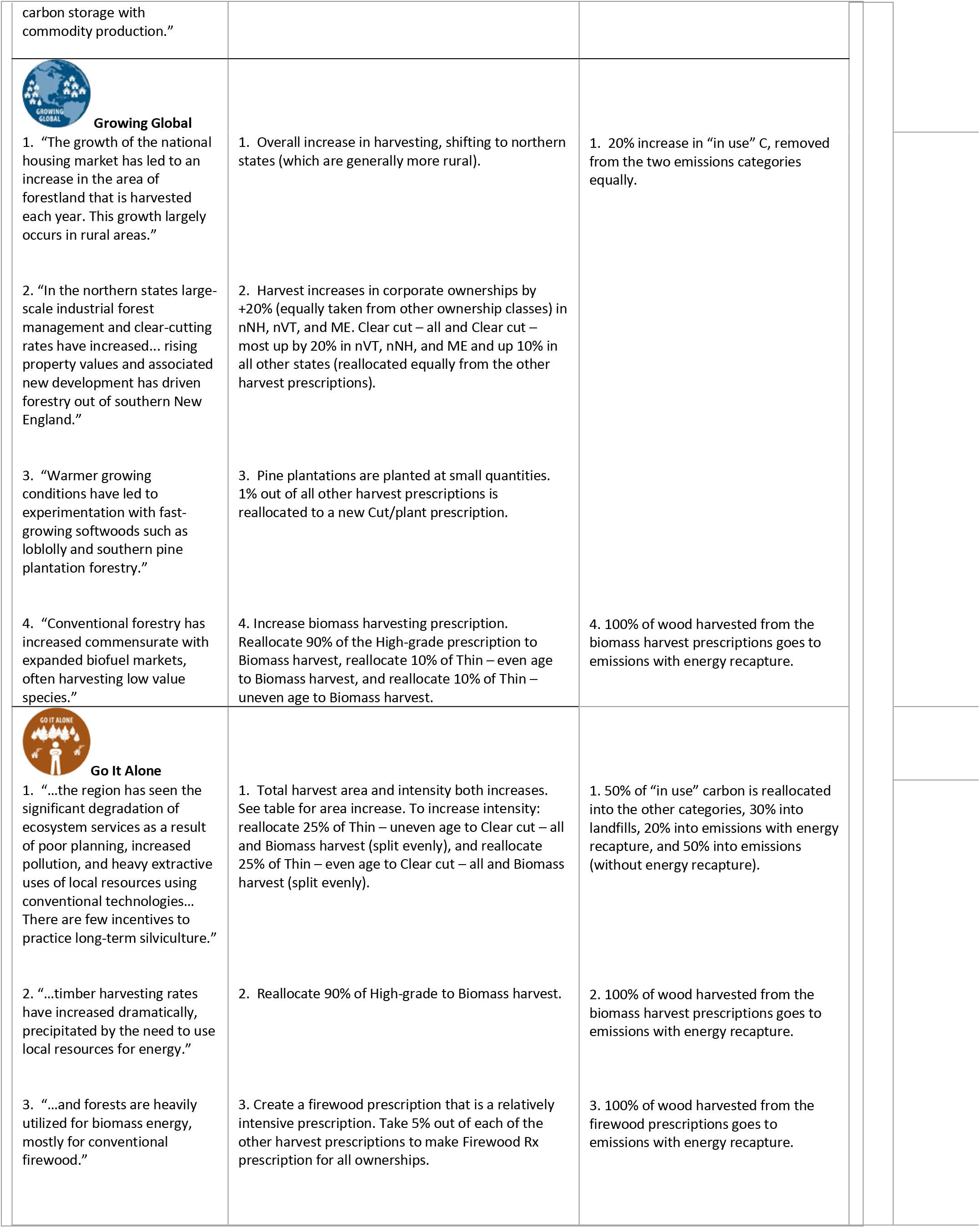
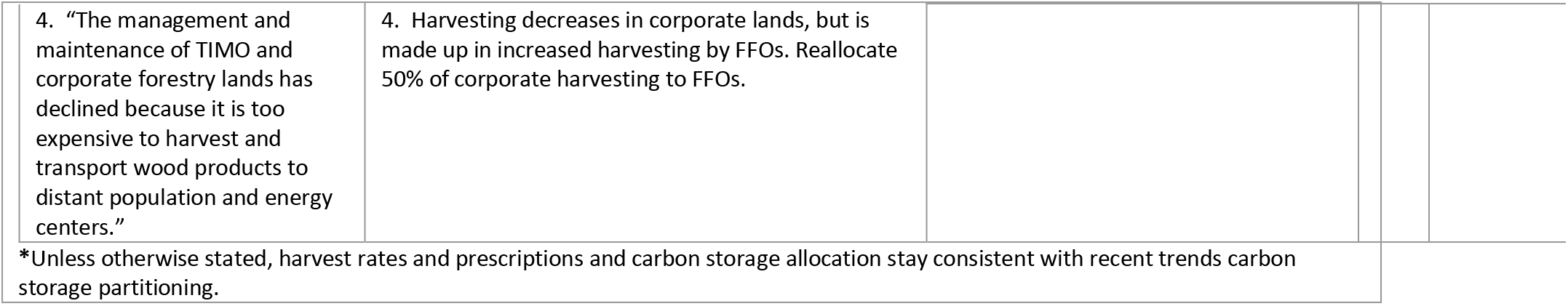
Representative quotes from stakeholder narratives and the implications for harvesting and carbon allocation. Most harvest implementations were given 40 years to ramp from the Recent Trends harvest rates to the envisioned 2060 harvest rates. Unless noted otherwise, changes to carbon allocation into new pools were implemented in year 10 of the simulation and then static for the remainder of the 50 years.

| Table 2. New harvesting prescriptions and carbon allocation |  |  |
| --- | --- | --- |
| Narrative Quotes<br>(Stakeholders) | Harvest Implementation | Carbon Allocation |
| **Connected Communities**<br>1. "...the use and protection of local resources increasingly important to governments and communities... there is a resurgence in community forests and woodlots near towns that are dedicated to producing high-value local wood products."<br>2. "A regional carbon tax... helps to promote greater | 1. Overall improvement in forest management. 60% reduction in rates of Clear cut - all and 40% reduction in High-grade and reallocated 80% to Ecological harvest – intense and 20% to Ecological harvest – light.<br>2. A Biomass Harvest prescription is added with a pulse of the Biomass Harvest prescription | 1. There is a 40% reduction in emitted with and without energy recaptures, as well as a 20% reduction in landfilling of carbon. The remaining carbon is "in use." |
| reliance on local food, local wood products and local transportation options during the early 2020s and 2030s, with local wood biomass serving as a renewable transition fuel.”<br>3. “...timber harvesting rates across the region increase by 50% by 2060, particularly in the northern New England states.” | implemented during timesteps 10 and 20, additional to the baseline harvest rates. The pulse is implemented by multiplying baseline Biomass Harvest rates by 1.5 in times 10 and 20.<br>3. Regional increase in total harvested area to 150% of RT in timestep 50. Increase more in the north than the south. | 2. 100% of wood harvested from the biomass harvest prescriptions goes to emissions with energy recapture. |
| **Yankee Cosmopolitan**<br>1. “Abundant forests remain a central part of New England’s identity, and support increases in tourism, particularly in Vermont, Maine, and New Hampshire... and carbon storage by forests is now highly valued.”<br>2. “Rates of timber harvesting for wood products have decreased in the region, particularly in southern New England where parcelization and non-timber forest values drive land management priorities.”<br>3. “Technological innovations in energy generation and storage limit the demand for wood biomass energy.”<br>4. “Development of sugar bushes has expanded as maple syrup has become a valuable global commodity and New England remains suitable for sugar maple trees despite changing climate.”<br>5. “...forestry practice laws designed to protect a range of ecosystem services have become more stringent in all states and the limited harvesting that occurs follows an ‘ecological forestry’ paradigm, including longer rotations with more leave trees and slash left on-site to balance | 1. Overall reduction in harvesting across New England.<br>2. Reduce overall harvesting, slightly more in southern NE than northern states. See table A5 for exact rates.<br>4. Take 1% out of each of the other harvest prescriptions to make a Maple Sugar prescription.<br>5. Rapid increase in Ecological Harvest prescriptions across NE. 90% of remaining harvest (after the sugarbush allocation) reallocated, 30% into Ecological Harvesting – intense and 70% into Ecological Harvesting – light by 2040 (ramped evenly). | 3. By 2060, 2/3 of the emissions do not have energy recapture. Linear replacement of emitted with energy recapture to emitted without energy recapture.<br>5. Slash left on site increases by 50%. There is a 20% reduction in both types of emissions, with a 10% reduction in landfilling of carbon. Remaining carbon is “in use”. |
| carbon storage with commodity production.” |  |  |
| **Growing Global**<br>1. “The growth of the national housing market has led to an increase in the area of forestland that is harvested each year. This growth largely occurs in rural areas.”<br>2. “In the northern states large-scale industrial forest management and clear-cutting rates have increased... rising property values and associated new development has driven forestry out of southern New England.”<br>3. “Warmer growing conditions have led to experimentation with fast-growing softwoods such as loblolly and southern pine plantation forestry.”<br>4. “Conventional forestry has increased commensurate with expanded biofuel markets, often harvesting low value species.” | 1. Overall increase in harvesting, shifting to northern states (which are generally more rural).<br>2. Harvest increases in corporate ownerships by +20% (equally taken from other ownership classes) in nNH, nVT, and ME. Clear cut – all and Clear cut – most up by 20% in nVT, nNH, and ME and up 10% in all other states (reallocated equally from the other harvest prescriptions).<br>3. Pine plantations are planted at small quantities. 1% out of all other harvest prescriptions is reallocated to a new Cut/plant prescription.<br>4. Increase biomass harvesting prescription. Reallocate 90% of the High-grade prescription to Biomass harvest, reallocate 10% of Thin – even age to Biomass harvest, and reallocate 10% of Thin – uneven age to Biomass harvest. | 1. 20% increase in “in use” C, removed from the two emissions categories equally.<br>4. 100% of wood harvested from the biomass harvest prescriptions goes to emissions with energy recapture. |
| **Go It Alone**<br>1. “...the region has seen the significant degradation of ecosystem services as a result of poor planning, increased pollution, and heavy extractive uses of local resources using conventional technologies... There are few incentives to practice long-term silviculture.”<br>2. “...timber harvesting rates have increased dramatically, precipitated by the need to use local resources for energy.”<br>3. “...and forests are heavily utilized for biomass energy, mostly for conventional firewood.” | 1. Total harvest area and intensity both increases. See table for area increase. To increase intensity: reallocate 25% of Thin – uneven age to Clear cut – all and Biomass harvest (split evenly), and reallocate 25% of Thin – even age to Clear cut – all and Biomass harvest (split evenly).<br>2. Reallocate 90% of High-grade to Biomass harvest.<br>3. Create a firewood prescription that is a relatively intensive prescription. Take 5% out of each of the other harvest prescriptions to make Firewood Rx prescription for all ownerships. | 1. 50% of “in use” carbon is reallocated into the other categories, 30% into landfills, 20% into emissions with energy recapture, and 50% into emissions (without energy recapture).<br>2. 100% of wood harvested from the biomass harvest prescriptions goes to emissions with energy recapture.<br>3. 100% of wood harvested from the firewood prescriptions goes to emissions with energy recapture. |

|  |  |
| --- | --- |
| 4. "The management and maintenance of TIMO and corporate forestry lands has declined because it is too expensive to harvest and transport wood products to distant population and energy centers." | 4. Harvesting decreases in corporate lands, but is made up in increased harvesting by FFOs. Reallocate 50% of corporate harvesting to FFOs. |
| *Unless otherwise stated, harvest rates and prescriptions and carbon storage allocation stay consistent with recent trends carbon storage partitioning. |  |

## APPENDIX III Harvest rates by management area and ownership class

Initial harvest rates were calculated using remeasured FIA plots within each of our management areas. Management areas were first defined by location/region and then ownership class, where regions were defined as states, with the exception of New Hampshire and Vermont, where the FIA definitions of northern and southern parts of the state were used since harvest regimes were different enough to warrant separate analyses. Next we simplified the FIA ownership classes (Table A3) to calculate the annual probability of harvest and harvest percent within each management area (as defined by region and ownership class), following methods used in Thompson et al. (2017a). The annual probability of harvest for each management area was calculated using the proportion of plots harvested, according to the FIA database, in the last three measurement periods (approx. 2000-2018) and the years between remeasurements to calculate an annual probability of harvest within each region and ownership class. We then calculated the average intensity of harvest by calculating the percent of the total biomass removed for those plots with a harvest within each management area. We then combined the annual probabilities and intensities of harvest in management areas with too few FIA plots (<100, with the exception of FFOs in RI), to the geographically nearest neighboring management area of the same ownership type with the most similar average harvesting probability and intensity (Table A4).

**Table A3.** Crosswalk between FIA ownership classes to our simplified ownerships for creating management areas. Note that final Management Areas for LANDIS-II included conservation status as well.

| OWNCD | FIA ownership class | Management areas | Name |
| --- | --- | --- | --- |
| 41 | corporate | CO | Corporate |
| 42 | NGO, natural resources organization | FF | Family Forest |
| 43 | unincorporated local partnership/club | LO | Local |
| 44 | Native American | FF | Family Forest |
| 45 | Individual | FF | Family Forest |
| 11 | National Forest | FE | Federal |
| 12 | National Grassland and/or Prairie | FE | Federal |
| 13 | Other Forest Service Land | FE | Federal |
| 21 | National Park Service | FE | Federal |
| 22 | Bureau of Land Management | FE | Federal |
| 23 | Fish and Wildlife Service | FE | Federal |
| 24 | Departments of Defense/Energy | FE | Federal |
| 25 | Other Federal | FE | Federal |
| 31 | State including State public universities | ST | State |
| 32 | Local (County, Municipality, etc.) including water authorities | LO | Local |
| 33 | Other Non-federal Public | LO | Local |
| 46 | Undifferentiated private | FF | Family Forest |

**Table A4.** FIA harvest intensities by management area Corporate owned forests

| Region(s) | # of FIA plots (n) | Annual Probability of Harvest | Average Harvest Intensity |
| --- | --- | --- | --- |
| ME | 1953 | 3.5% | 45.1% |
| nNH, nVT | 160 | 3.5% | 47.0% |
| sNH, sVT | 121 | 1.9% | 24.3% |
| CT, MA, RI | 145 | 0.92% | 28.5% |

Family owned forests
| Region(s) | # of FIA plots (n) | Annual Probability of Harvest | Average Harvest Intensity |
| --- | --- | --- | --- |
| ME | 1327 | 3.7% | 29.0% |
| nNH | 204 | 2.8% | 28.1% |
| sNH | 322 | 2.5% | 14.2% |
| nVT | 330 | 2.8% | 28.5% |
| sVT | 265 | 2.6% | 22.8% |
| MA | 337 | 1.6% | 17.0% |
| CT | 225 | 2.4% | 11.2% |
| RI | 88 | 1.1% | 19.0% |

Federally owned forests
| Region(s) | # of FIA plots (n) | Annual Probability of Harvest | Average Harvest Intensity |
| --- | --- | --- | --- |
| All regions together | 363 | 0.61% | 37.4% |

Locally owned forests
| Region(s) | # of FIA plots (n) | Annual Probability of Harvest | Average Harvest Intensity |
| --- | --- | --- | --- |
| ME, nNH, sNH, nVT, sVT | 211 | 3.0% | 18.0% |
| CT, MA, RI | 181 | 1.2% | 19.6% |

State owned forests
| Region(s) | # of FIA plots (n) | Annual Probability of Harvest | Average Harvest Intensity |
| --- | --- | --- | --- |
| ME | 139 | 1.8% | 35.5% |
| **nNH, sNH, nVT,**<br>**sVT** | 102 | 0.78% | 30.7% |
| **CT, MA, RI** | 174 | 1.0% | 31.8% |

These calculated annual probabilities of harvest were used as the harvest rates for each management area for the Recent Trends (RT) simulation. The average harvest intensities for each management area were used as the target average intensity of harvest for each management area. A linear programming method was used to balance the individual intensities of each of the harvest prescriptions so that the average harvest intensity for each management area was within 1% of the target average harvest intensity. Initial allocation of harvest proportions was based on the work of Belair and Ducey (2018), from which the linear programming method rebalanced the allocation to meet some given requirements and the target average harvest intensity. The harvest allocation models met the following requirements: (1) all harvest proportions together must be 100% of harvests for that management area; (2) different prescriptions could vary more than others (Clear cut – all (+20%, −10%), Thin – uneven age (±20%), all others (±5%)), no harvest types could go to zero (lowest proportion = 0.1%), and no harvest types could go to 100%. All models converged.

Finally, the individual scenario narratives were used to alter both the harvest rates and intensities for each of the divergent scenarios. First, the overall target harvest area was either increased or decreased for each management area (Table A5) and then translated into new rates given the available area for harvest in each management area. Next, scenario descriptions were used to reallocate harvests in RT to different and/or newly defined harvest prescriptions specific to each scenario (Table A6).

**Table A5.** Target harvested area as a percent of area harvested in the RT scenario.

| **Scenario** | **Region** | **Overall harvest area in 2060**<br>**as a % of RT** |
| --- | --- | --- |
| **YC** | ME, nNH, sNH, nVT, sVT<br>MA, CT, RI | 60%<br>40% |
| **GG** | ME<br>nNH, nVT<br>sNH, sVT<br>MA, CT, RI | 250%<br>200%<br>150%<br>50% |
| **GA** | All management areas | 250% |
| **CC** | ME<br>nNH, sNH, nVT, sVT<br>MA, CT, RI | 165%<br>140%<br>115% |

**Table A6.** Target prescription allocations in the final year (2060)

| **Rx (weighted by**<br>**area across all**<br>**management**<br>**areas)** | **RT year 50**<br>**(10 year %**<br>**harv)** | **CC year 50**<br>**(10 year %**<br>**harv)** | **YC year 50**<br>**(10 year %**<br>**harv)** | **GG year 50**<br>**(10 year %**<br>**harv)** | **GA year 50**<br>**(10 year %**<br>**harv)** |
| --- | --- | --- | --- | --- | --- |
| **Clear cut - all** | 2.73% | 0.81% | 0.16 % | 8.41% | 10.50% |
| **Clear cut - most** | 1.87% | 0.56% | 0.11% | 6.52% | 2.28% |
| **High-grade** | 3.57% | 0.32% | 0.20% | 0.72% | 0.93% |
| **Shelterwood** | 2.62% | 3.92% | 0.15% | 5.37% | 5.61% |
| **Thin – even age** | 5.77% | 8.87% | 0.33% | 11.90% | 8.42% |
| **Thin - uneven age** | 11.83% | 17.83% | 0.67% | 22.34% | 20.67% |
| **Biomass harvest** | 0.00% | 4.25% | 0.00% | 10.31% | 13.20% |
| **Cut/plant** | 0.00% | 0.00% | 0.00% | 0.66% | 0.00% |
| **Ecological harvest – intense** | 0.00% | 5.73% | 4.41% | 0.00% | 0.00% |
| **Ecological harvest - light** | 0.00% | 1.52% | 10.29% | 0.00% | 0.00% |
| **Firewood cutting** | 0.00% | 0.00% | 0.00% | 0.00% | 3.24% |
| **Sugarbush** | 0.00% | 0.00% | 0.17% | 0.00% | 0.00% |

## APPENDIX IV Carbon allocation process

Our harvested carbon accounting framework resulted in estimates of carbon emitted through decomposition or combustion, emitted with energy recapture (e.g., used in energy generation), still in use (e.g., in wood product such as building material), landfilled, and still in slash (not decomposed yet) for the harvested carbon for the entire simulation. We only tracked the carbon impacted by harvest during our simulation time period, from 2010-2060; therefore, any carbon stored or emitted as a result of harvesting previous to 2010, or transitions that happened after 2060 (e.g., from in-use to emitted), were not included in our accounting.

Specifically, to partition removed growing stock carbon (GSC_R_; after slash removal) into saw timber and pole timber, the forest type and hardwood/softwood specific values from Smith et al., (2006; Table 4) were used in accordance with the following:

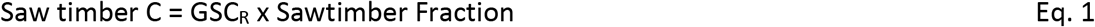

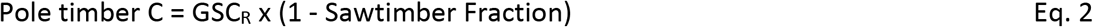

Next, the appropriate values from Smith et al., (2006; Table 5) were used to partition the saw timber and pole timber into saw log, pulp wood, bark and fuel wood using the following (also with values specific to wood type):

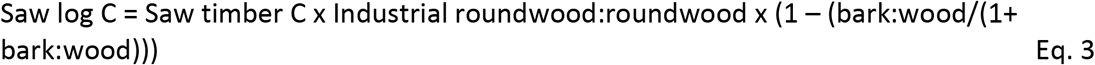

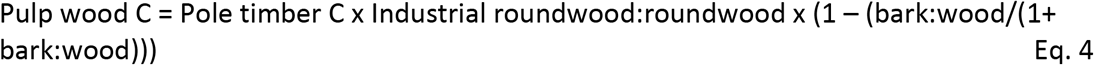

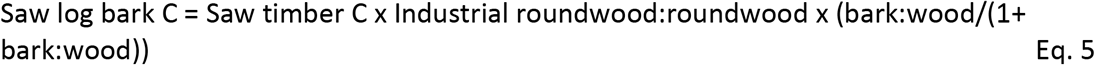

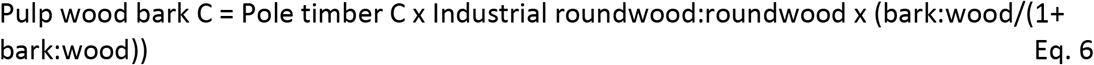

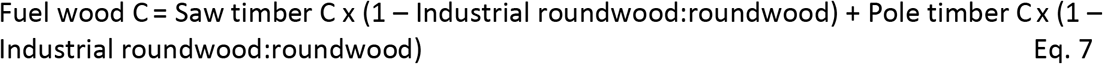

Finally, decay rates for slash (Russell et al. 2014) and bark tables (Smith et al. 2006) were used to allocate the removed wood to the final tracked carbon pools by time since removal, using the following:

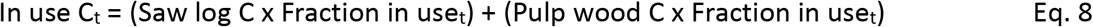

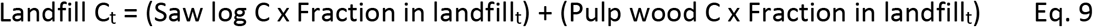

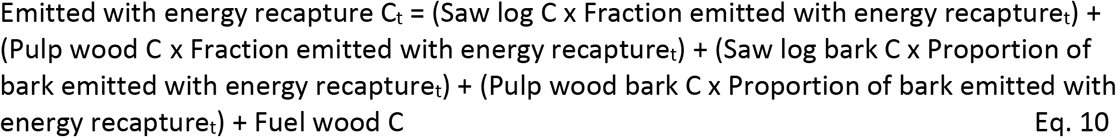

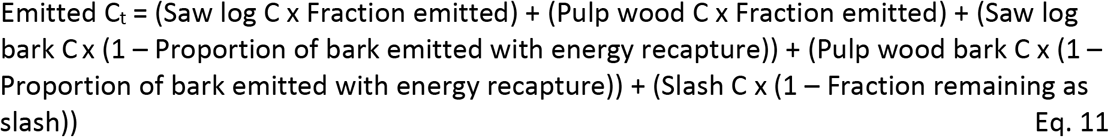

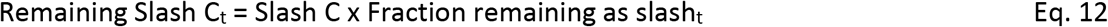

Where *t* is the fraction allocated to each pool specific to the time since harvest. For example, as time since harvest increases, the amount of the total removed carbon that is “in use” decreases while the amount that is “emitted” or “landfilled” increases.

## APPENDIX V Removed carbon allocation by carbon removal type by scenario

Each scenario had different carbon removal processes at play, resulting in different contributions to both emissions and storage pools. Below is the breakdown of the removed carbon in pools by time step, removal type, and scenario.

**Figure A1.**
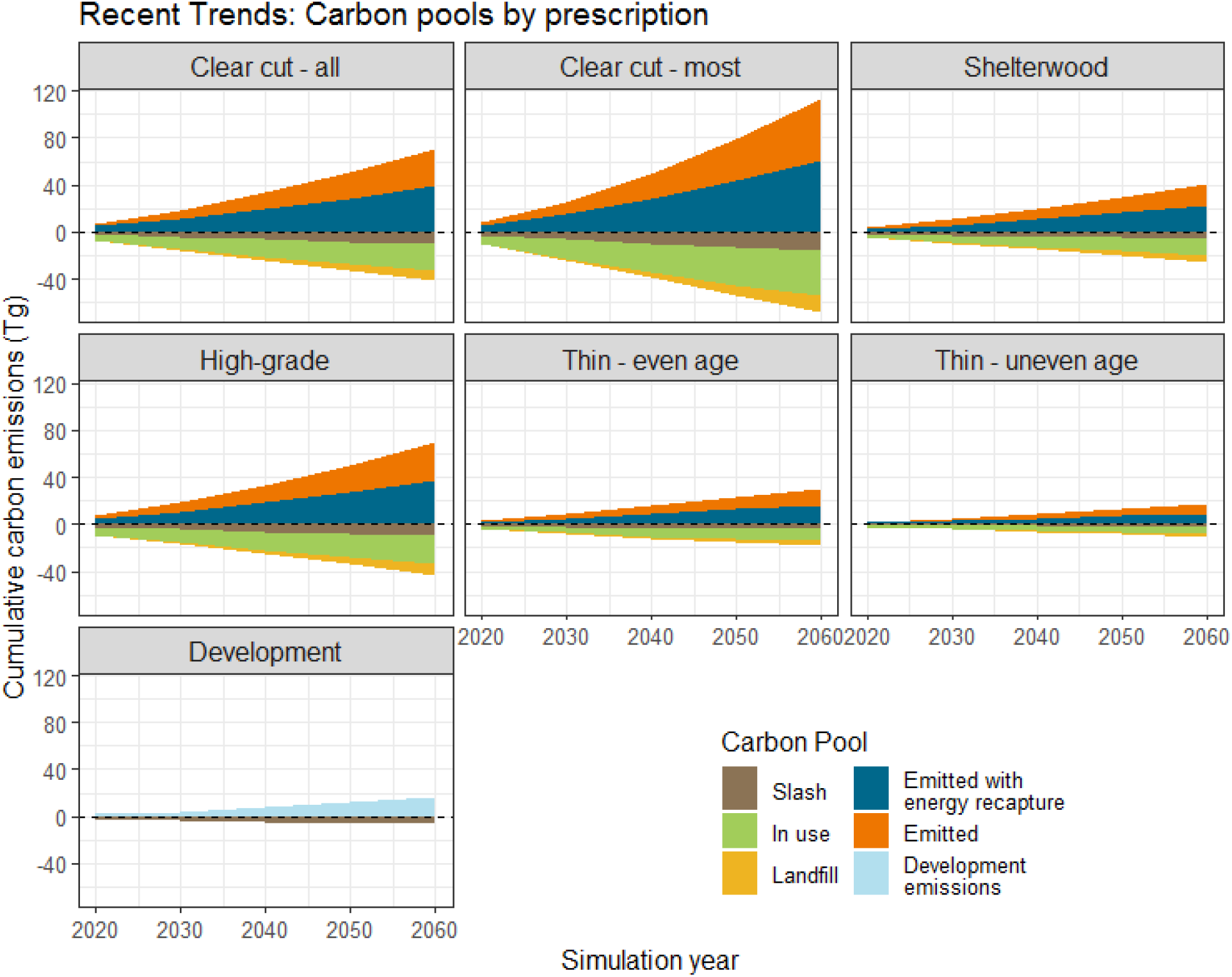
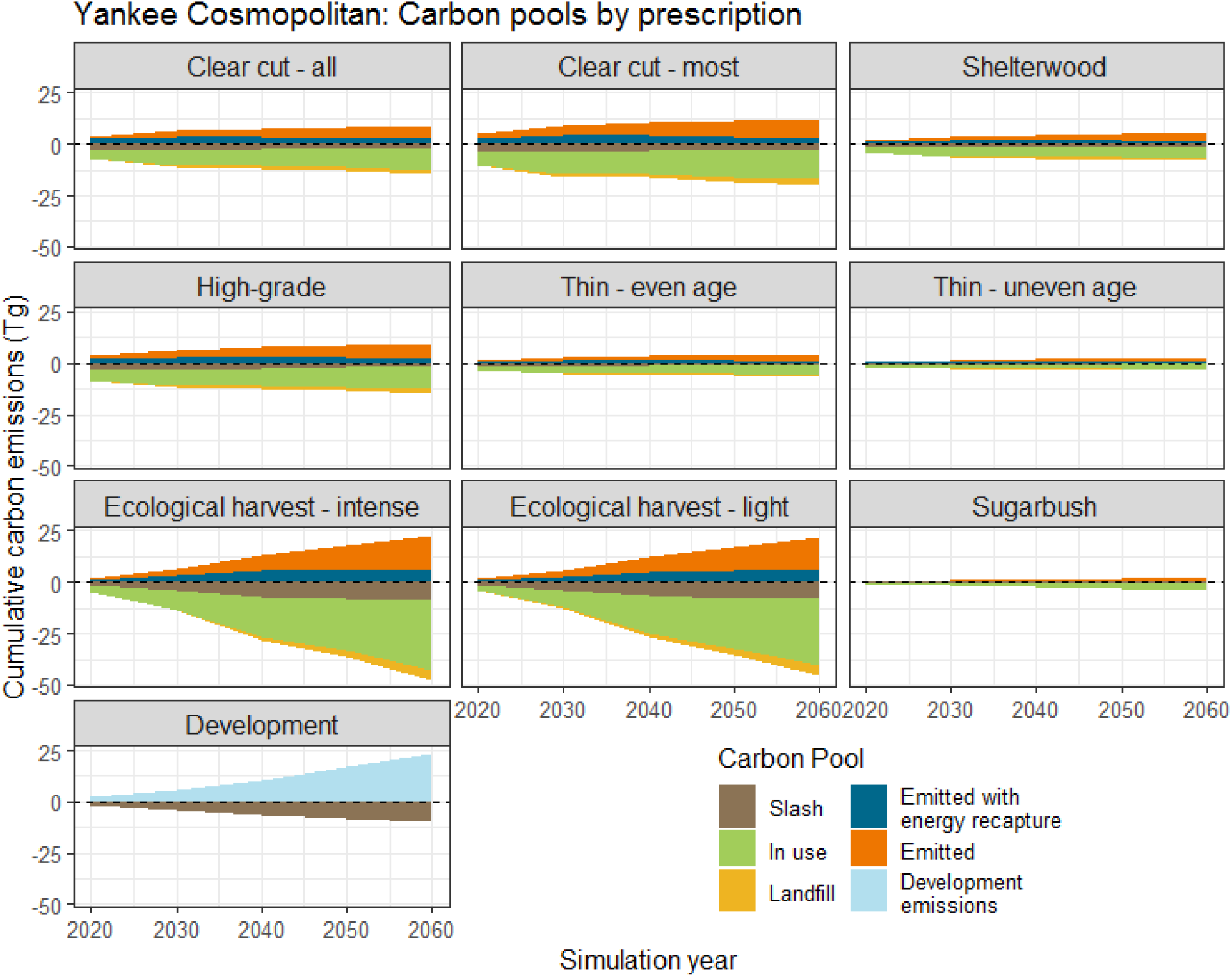
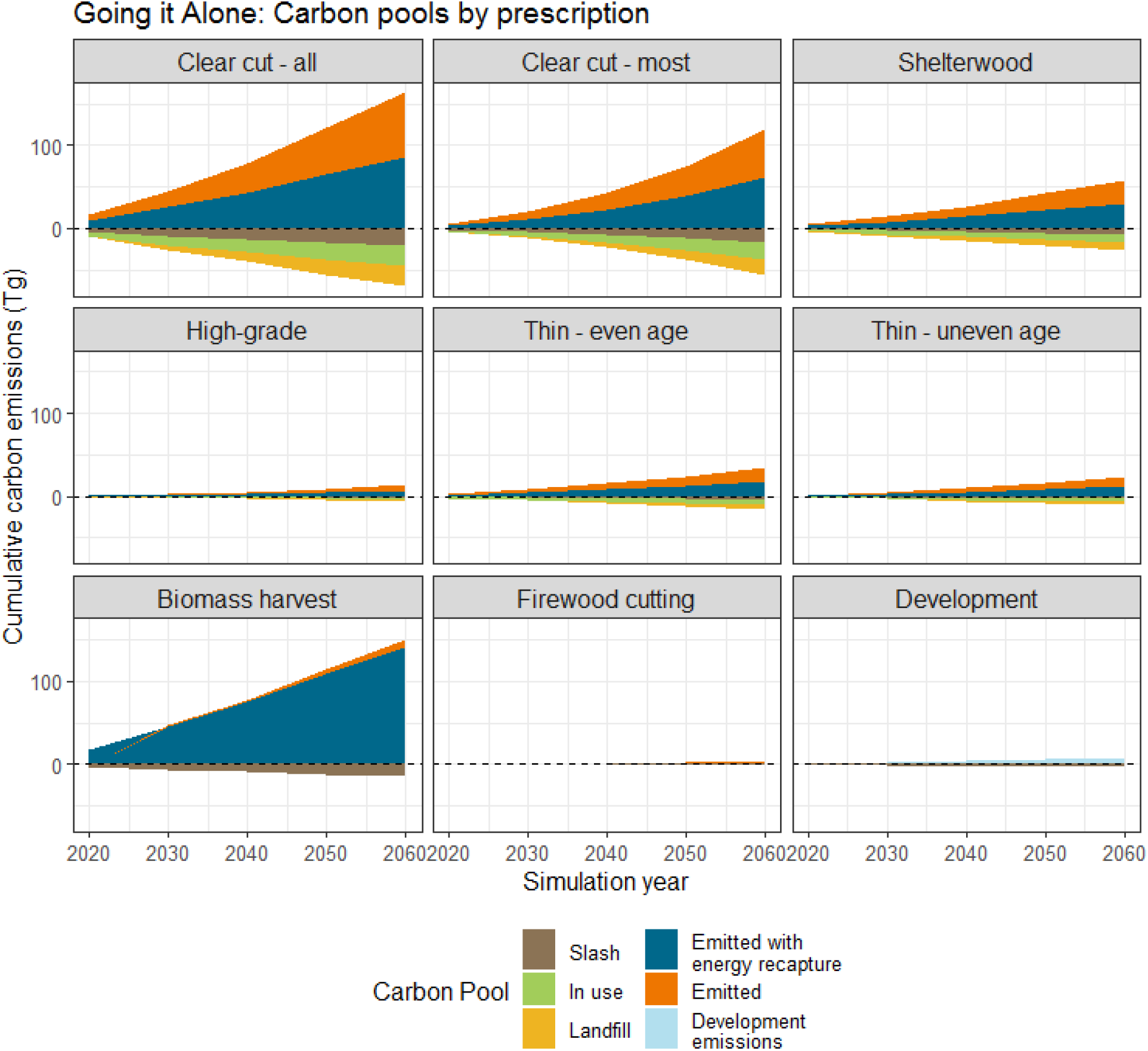
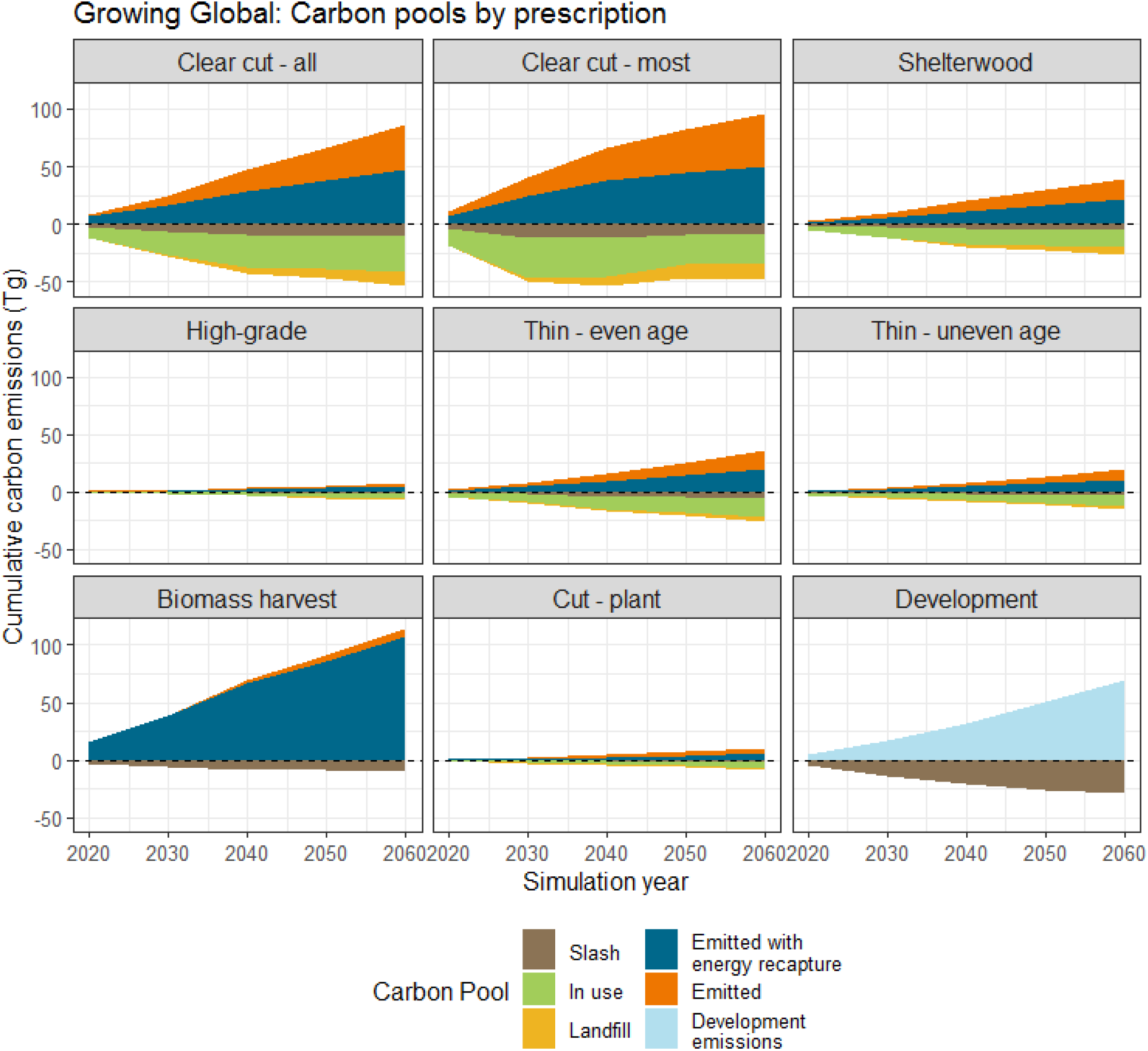
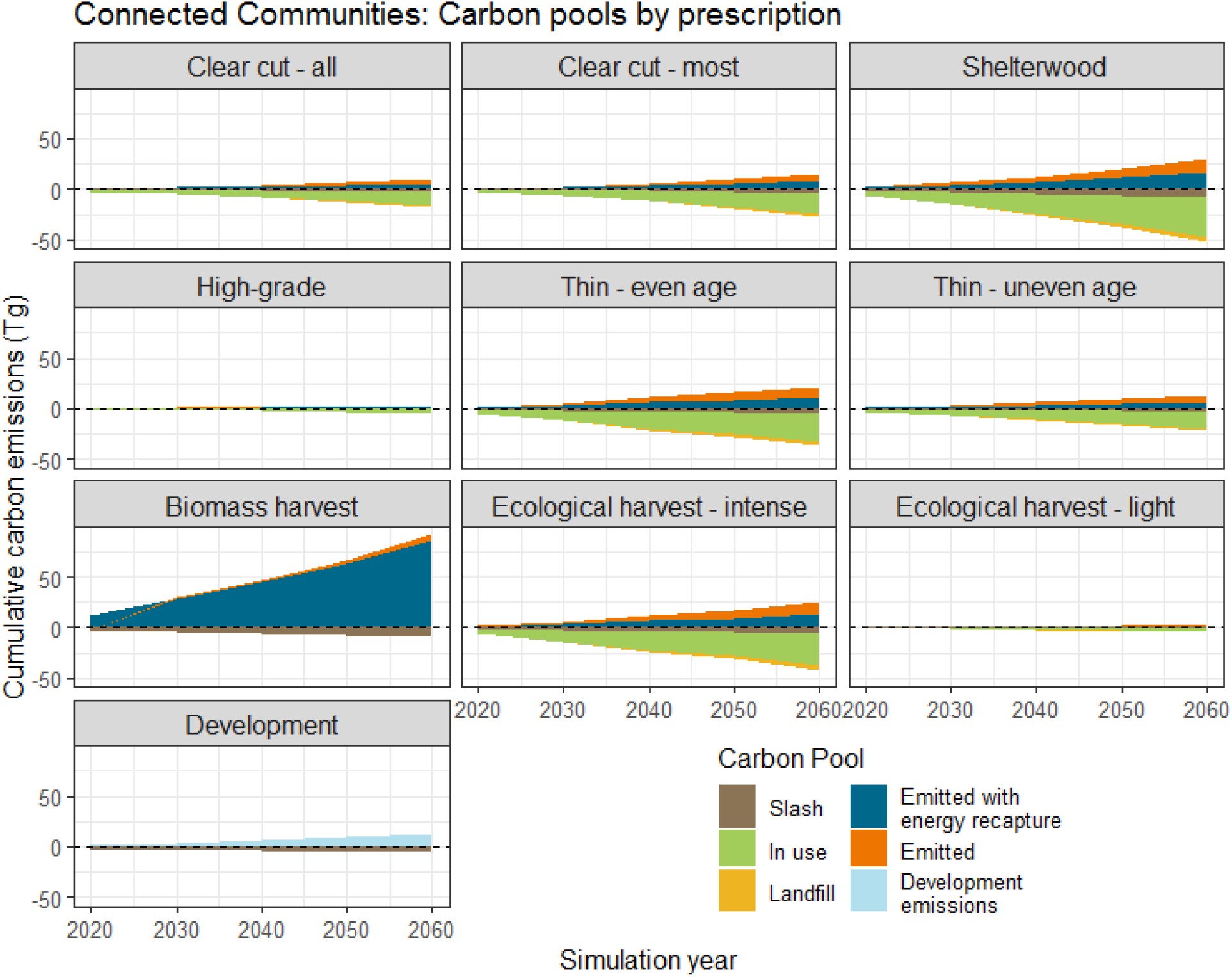
Removed carbon pools by removal type and scenario.

## CITATIONS

Aber, J. D., S. V. Ollinger, C. A. Federer, P. B. Reich, M. L. Goulden, D. W. Kicklighter, J. M. Melillo, and R. G. Lathrop. 1995. Predicting the effects of climate change on water yield and forest production in the northeastern United States. Climate Research 5:207–222.

Bechtold, W. A., and P. L. Patterson. 2005. The Enhanced Forest Inventory and Analysis Program-National Sampling Design and Estimation Procedures. U.S. Department of Agriculture, Forest Service, Southern Research Station, Ashville, NC.

Belair, E. P., and M. J. Ducey. 2018. Patterns in Forest Harvesting in New England and New York: Using FIA Data to Evaluate Silvicultural Outcomes. Journal of Forestry 116:273–282.

Bradfield, R., G. Wright, G. Burt, G. Cairns, and K. Van Der Heijden. 2005. The origins and evolution of scenario techniques in long range business planning.

de Bruijn, A., E. J. Gustafson, B. R. Sturtevant, J. R. Foster, B. R. Miranda, N. I. Lichti, and D. F. Jacobs. 2014. Toward more robust projections of forest landscape dynamics under novel environmental conditions: Embedding PnET within LANDIS-II. Ecological Modelling 287:44–57.

Butler, B. J., S. J. Crocker, G. M. Domke, C. M. Kurtz, T. W. Lister, P. D. Miles, R. S. Morin, R. J. Piva, R. Riemann, and C. W. Woodall. 2015. The forests of Southern New England, 2012.

Butler, B. J., J. H. Hewes, B. J. Dickinson, K. Andrejczyk, S. M. Butler, and M. Markowski-Lindsay. 2016. Family Forest Ownerships of the United States, 2013: Findings from the USDA Forest Service’s National Woodland Owner Survey. Journal of Forestry 114:638–647.

Canham, C. D., N. Rogers, and T. Buchholz. 2013. Regional variation in forest harvest regimes in the northeastern United States. Ecological Applications 23:515–522.

Cook-Patton, S. C., S. M. Leavitt, D. Gibbs, N. L. Harris, K. Lister, K. J. Anderson-Teixeira, R. D. Briggs, R. L. Chazdon, T. W. Crowther, P. W. Ellis, H. P. Griscom, V. Herrmann, K. D. Holl, R. A. Houghton, C. Larrosa, G. Lomax, R. Lucas, P. Madsen, Y. Malhi, A. Paquette, J. D. Parker, K. Paul, D. Routh, S. Roxburgh, S. Saatchi, J. van den Hoogen, W. S. Walker, C. E. Wheeler, S. A. Wood, L. Xu, and B. W. Griscom. 2020a. Mapping carbon accumulation potential from global natural forest regrowth. Nature 585:545–550.

Cook-Patton, S. C., S. M. Leavitt, D. Gibbs, N. L. Harris, K. Lister, K. J. Anderson-Teixeira, R. D. Briggs, R. L. Chazdon, T. W. Crowther, P. W. Ellis, H. P. Griscom, V. Herrmann, K. D. Holl, R. A. Houghton, C. Larrosa, G. Lomax, R. Lucas, P. Madsen, Y. Malhi, A. Paquette, J. D. Parker, K. Paul, D. Routh, S. Roxburgh, S. Saatchi, J. van den Hoogen, W. S. Walker, C. E. Wheeler, S. A. Wood, L. Xu, and B. W. Griscom. 2020b. Mapping carbon accumulation potential from global natural forest regrowth. Nature 585:545–550.

Duveneck, M. J., and J. R. Thompson. 2017. Climate change imposes phenological trade-offs on forest net primary productivity. Journal of Geophysical Research: Biogeosciences 122:2298–2313.

Duveneck, M. J., and J. R. Thompson. 2019. Social and biophysical determinants of future forest conditions in New England: Effects of a modern land-use regime. Global Environmental Change 55:115–129.

Duveneck, M. J., J. R. Thompson, E. J. Gustafson, Y. Liang, and A. M. G. de Bruijn. 2017. Recovery dynamics and climate change effects to future New England forests. Landscape Ecology 32:1385–1397.

Duveneck, M. J., J. R. Thompson, and B. T. Wilson. 2015. An imputed forest composition map for New England screened by species range boundaries. Forest Ecology and Management 347:107–115.

FHWA. 2015. Office of Highway Policy Information - Policy | Federal Highway Administration. https://www.fhwa.dot.gov/policyinformation/statistics/2010/mv1.cfm.

Finzi, A. C., M. Giasson, A. A. Barker Plotkin, J. D. Aber, E. R. Boose, E. A. Davidson, M. C. Dietze, A. M. Ellison, S. D. Frey, E. Goldman, T. F. Keenan, J. M. Melillo, J. W. Munger, K. J. Nadelhoffer, S. V. Ollinger, D. A. Orwig, N. Pederson, A. D. Richardson, K. Savage, J. Tang, J. R. Thompson, C. A. Williams, S. C. Wofsy, Z. Zhou, and D. R. Foster. 2020. Carbon budget of the Harvard Forest Long-Term Ecological Research site: pattern, process, and response to global change. Ecological Monographs:ecm.1423.

Gunn, J. S., M. J. Ducey, and E. Belair. 2019. Evaluating degradation in a North American temperate forest. Forest Ecology and Management 432:415–426.

Gustafson, E. J. 2013. When relationships estimated in the past cannot be used to predict the future: using mechanistic models to predict landscape ecological dynamics in a changing world. Landscape Ecology 28:1429–1437.

Gustafson, E. J., S. R. Shifley, D. J. Mladenoff, K. K. Nimerfro, and H. S. He. 2000. Spatial simulation of forest succession and timber harvesting using LANDIS. Canadian Journal of Forest Research 30:32–43.

Guswa, A. J., B. Hall, C. Cheng, and J. R. Thompson. 2020. Co-designed Land-use Scenarios and their Implications for Storm Runoff and Streamflow in New England. Environmental Management:1–16.

Harris, N. L., S. C. Hagen, S. S. Saatchi, T. R. H. Pearson, C. W. Woodall, G. M. Domke, B. H. Braswell, B. F. Walters, S. Brown, W. Salas, A. Fore, and Y. Yu. 2016. Attribution of net carbon change by disturbance type across forest lands of the conterminous United States. Carbon Balance Manage 11:24.

Henders, S., and M. Ostwald. 2012. Forest Carbon Leakage Quantification Methods and Their Suitability for Assessing Leakage in REDD. Forests 3:33–58.

Houghton, R. A., J. I. House, J. Pongratz, G. R. Van Der Werf, R. S. Defries, M. C. Hansen, C. Le Quéré, and N. Ramankutty. 2012. Carbon emissions from land use and land-cover change. Biogeosciences 9:5125–5142.

Huntington, T. G., A. D. Richardson, K. J. McGuire, and K. Hayhoe. 2009. Climate and hydrological changes in the northeastern United States: Recent trends and implications for forested and aquatic ecosystems. Canadian Journal of Forest Research 39:199–212.

Jevon, F. V., A. W. D’Amato, C. W. Woodall, K. Evans, M. P. Ayres, and J. H. Matthes. 2019. Tree basal area and conifer abundance predict soil carbon stocks and concentrations in an actively managed forest of northern New Hampshire, USA. Forest Ecology and Management 451:117534.

Kaboli, H., P. L. Clouston, and S. Lawrence. 2020. Feasibility of Two Northeastern Species in Three-Layer ANS1-Approved Cross-Laminated Timber. Journal of Materials in Civil Engineering 32:04020006.

Kittredge, D. B., J. R. Thompson, L. L. Morreale, A. G. Short Gianotti, and L. R. Hutyra. 2017. Three decades of forest harvesting along a suburban-rural continuum. Ecosphere 8d:e01882.

Li, H., Y. Ma, T. M. Aide, and W. Liu. 2008. Past, present and future land-use in Xishuangbanna, China and the implications for carbon dynamics. Forest Ecology and Management 255:16–24.

Liang, Y., M. J. Duveneck, E. J. Gustafson, J. M. Serra-Diaz, and J. R. Thompson. 2018. How disturbance, competition, and dispersal interact to prevent tree range boundaries from keeping pace with climate change. Global Change Biology 24:e335–e351.

Losing Ground: Nature’s Value in a Changing Climate, Sixth Edition of the Losing Ground series. 2020.

Ma, W., G. M. Domke, C. W. Woodall, and A. W. D’Amato. 2020. Contemporary forest carbon dynamics in the northern U.S. associated with land cover changes. Ecological Indicators 110:105901.

McBride, M. F., M. J. Duveneck, K. F. Lambert, K. A. Theoharides, and J. R. Thompson. 2019. Perspectives of resource management professionals on the future of New England’s landscape: Challenges, barriers, and opportunities. Landscape and Urban Planning 188.

McBride, M. F., K. F. Lambert, E. S. Huff, K. A. Theoharides, P. Field, and J. R. Thompson. 2017. Increasing the effectiveness of participatory scenario development through codesign. Ecology and Society 22:art16.

McKenzie, P. F., M. J. Duveneck, L. L. Morreale, and J. R. Thompson. 2019. Local and global parameter sensitivity within an ecophysiologically based forest landscape model. Environmental Modelling and Software 117:1–13.

Mladenoff, D. J., and H. S. He. 1999. Design, behavior and application of LANDIS, an object-oriented model of forest landscape disturbance and succession. Spatial modeling of forest landscape change: approaches and applications. Papers presented at a symposium in Albuquerque, New Mexico, USA, 1997.:125–162.

New England Forestry Foundation. 2017. Assessing the wood supply and investment potential for a New England engineered wood products mill.

Olofsson, P., C. E. Holden, E. L. Bullock, and C. E. Woodcock. 2016. Time series analysis of satellite data reveals continuous deforestation of New England since the 1980s. Environmental Research Letters 11:064002.

Pan, Y., R. A. Birdsey, J. Fang, R. Houghton, P. E. Kauppi, W. A. Kurz, O. L. Phillips, A. Shvidenko, S. L. Lewis, J. G. Canadell, P. Ciais, R. B. Jackson, S. W. Pacala, A. D. McGuire, S. Piao, A. Rautiainen, S. Sitch, and D. Hayes. 2011. A large and persistent carbon sink in the world’s forests. Science 333:988–993.

Pearman-Gillman, S. B., M. J. Duveneck, J. D. Murdoch, and T. M. Donovan. 2020a. Drivers and Consequences of Alternative Landscape Futures on Wildlife Distributions in New England, United States. Frontiers in Ecology and Evolution 8:164.

Pearman-Gillman, S. B., M. J. Duveneck, J. D. Murdoch, and T. M. Donovan. 2020b. Wildlife resistance and protection in a changing New England landscape. PLOS ONE 15:e0239525.

Popp, A., F. Humpenöder, I. Weindl, B. L. Bodirsky, M. Bonsch, H. Lotze-Campen, C. Müller, A. Biewald, S. Rolinski, M. Stevanovic, and J. P. Dietrich. 2014. Land-use protection for climate change mitigation. Nature Climate Change 4:1095–1098.

Puhlick, J., C. Woodall, and A. Weiskittel. 2017. Implications of land-use change on forest carbon stocks in the eastern United States. Environmental Research Letters 12:024011.

Le Quéré, C., R. M. Andrew, P. Friedlingstein, S. Sitch, J. Pongratz, A. C. Manning, J. Ivar Korsbakken, G. P. Peters, J. G. Canadell, R. B. Jackson, T. A. Boden, P. P. Tans, O. D. Andrews, V. K. Arora, D. C. E. Bakker, L. Barbero, M. Becker, R. A. Betts, L. Bopp, F. Chevallier, L. P. Chini, P. Ciais, C. E. Cosca, J. Cross, K. Currie, T. Gasser, I. Harris, J. Hauck, V. Haverd, R. A. Houghton, C. W. Hunt, G. Hurtt, T. Ilyina, A. K. Jain, E. Kato, M. Kautz, R. F. Keeling, K. Klein Goldewijk, A. Körtzinger, P. Landschützer, N. Lefèvre, A. Lenton, S. Lienert, I. Lima, D. Lombardozzi, N. Metzl, F. Millero, P. M. S. Monteiro, D. R. Munro, J. E. M. S. Nabel, S. I. Nakaoka, Y. Nojiri, X. Antonio Padin, A. Peregon, B. Pfeil, D. Pierrot, B. Poulter, G. Rehder, J. Reimer, C. Rödenbeck, J. Schwinger, R. Séférian, I. Skjelvan, B. D. Stocker, H. Tian, B. Tilbrook, F. N. Tubiello, I. T. V. Laan-Luijkx, G. R. V. Werf, S. Van Heuven, N. Viovy, N. Vuichard, A. P. Walker, A. J. Watson, A. J. Wiltshire, S. Zaehle, and D. Zhu. 2018. Global Carbon Budget 2017. Earth System Science Data 10:405–448.

Reinmann, A. B., L. R. Hutyra, A. Trlica, and P. Olofsson. 2016. Assessing the global warming potential of human settlement expansion in a mesic temperate landscape from 2005 to 2050. Science of the Total Environment 545-546:512–524.

Russell, M. B., C. W. Woodall, S. Fraver, A. W. D’Amato, G. M. Domke, and K. E. Skog. 2014. Residence Times and Decay Rates of Downed Woody Debris Biomass/Carbon in Eastern US Forests. Ecosystems 17:765–777.

Scheller, R. M. 2020. LANDIS-II Biomass Community Output v2.0.

Scheller, R. M., J. B. Domingo, B. R. Sturtevant, J. S. Williams, A. Rudy, E. J. Gustafson, and D. J. Mladenoff. 2007. Design, development, and application of LANDIS-II, a spatial landscape simulation model with flexible temporal and spatial resolution. Ecological Modelling 201:409–419.

Schulp, C. J. E., G. J. Nabuurs, and P. H. Verburg. 2008. Future carbon sequestration in Europe-Effects of land use change. Agriculture, Ecosystems and Environment 127:251–264.

Sleeter, B. M., J. Liu, C. Daniel, B. Rayfield, J. Sherba, T. J. Hawbaker, Z. Zhu, P. C. Selmants, and T. R. Loveland. 2018. Effects of contemporary land-use and land-cover change on the carbon balance of terrestrial ecosystems in the United States. Environmental Research Letters 13:045006.

Sleeter, B. M., T. L. Sohl, M. A. Bouchard, R. R. Reker, C. E. Soulard, W. Acevedo, G. E. Griffith, R. R. Sleeter, R. F. Auch, K. L. Sayler, S. Prisley, and Z. Zhu. 2012. Scenarios of land use and land cover change in the conterminous United States: Utilizing the special report on emission scenarios at ecoregional scales. Global Environmental Change 22:896–914.

Smith, J. E., L. S. Heath, K. E. Skog, and R. A. Birdsey. 2006. Methods for Calculating Forest Ecosystem and Harvested Carbon with Standard Estimates for Forest Types of the United States. USDA Northern Research Station General Te:216.

Stocker, T. F., D. Qin, G.-K. Plattner, M. M. B. Tignor, S. K. Allen, J. Boschung, A. Nauels, Y. Xia, V. Bex, and P. M. Midgley. 2013. Climate Change 2013 The Physical Science Basis Working Group I Contribution to the Fifth Assessment Report of the Intergovernmental Panel on Climate Change Edited by.

Stoner, A. M. K., K. Hayhoe, X. Yang, and D. J. Wuebbles. 2013. An asynchronous regional regression model for statistical downscaling of daily climate variables. International Journal of Climatology 33:2473–2494.

Thompson, J. R., C. D. Canham, L. Morreale, D. B. Kittredge, and B. Butler. 2017a. Social and biophysical variation in regional timber harvest regimes. Ecological Applications 27:942–955.

Thompson, J. R., D. N. Carpenter, C. V. Cogbill, and D. R. Foster. 2013. Four Centuries of Change in Northeastern United States Forests. PLoS ONE 8:e72540.

Thompson, J. R., J. Plisinski, K. Fallon Lambert, M. J. Duveneck, L. Morreale, M. McBride, M. Graham MacLean, M. Weiss, and L. Lee. 2020. Spatial simulation of co-designed land-cover change scenarios in New England: Alternative futures and their consequences for conservation priorities. Earth’s Future:e2019EF001348.

Thompson, J. R., J. S. Plisinski, P. Olofsson, C. E. Holden, and M. J. Duveneck. 2017b. Forest loss in New England: A projection of recent trends. PLoS ONE 12.

Williams, C. A., G. J. Collatz, J. Masek, and S. N. Goward. 2012. Carbon consequences of forest disturbance and recovery across the conterminous United States. Global Biogeochemical Cycles 26:n/a-n/a.

Woodall, C. W., J. W. Coulston, G. M. Domke, B. F. Walters, D. N. Wear, J. E. Smith, H. E. Andersen, B. J. Clough, W. B. Cohen, D. M. Griffith, S. C. Hagen, I. S. Hanou, M. C. Nichols, C. H. Perry, M. B. Russell, J. A. Westfall, B. T. Wilson, Woodall, Christopher W, Coulston, John W, Domke, Grant M, Walters, Brian F, Wear, David N, Smith, James E, Andersen, Hans-Erik, Clough, Brian J, Cohen, Warren B, Griffith, Douglas M, Hagen, Stephen C, Hanou, Ian S, Nichols, Michael C, Perry, Charles H, Russell, Matthew B, Westfall, James A, Wilson, and Barry T. 2015. The U.S. Forest Carbon Accounting Framework: Stocks and Stock Change, 1990-2016. Gen. Tech. Rep. NRS-154. Newtown Square, PA: U.S. Department of Agriculture, Forest Service, Northern Research Station. 49 p. 154:1–49.

Zhu, Z., and C. E. Woodcock. 2014. Continuous change detection and classification of land cover using all available Landsat data. Remote Sensing of Environment 144:152–171.

